# Evolution of mental traits linked to cultural development in modern humans and Neanderthals

**DOI:** 10.1101/2024.08.13.604521

**Authors:** Yuka Moriya, Naoko T. Fujito, Masahiro Terahara, Naoyuki Takahata, Yoko Satta, Kosuke M. Teshima, Toshiyuki Hayakawa

## Abstract

A cumulative process of cultural evolution occurred globally alongside the spread of anatomically modern humans (AMHs). This process of evolution was likely accompanied by global changes in mental and behavioral phenotypes, resulting in cultural differences between AMHs and Neanderthals. Globally, selective increases of frequencies were detected in alleles, which contributed to the suppression of neuroticism and advanced learning. This finding indicates that neuroticism was suppressed and learning behaviors developed through the global spread of AMHs. It suggests that global adaptation to psychosocial stress occurred, which contributed to cumulative cultural evolution by increasing cooperative learning. A comparison of phenotypic trends at the population level indicates that such globally occurring adaptive evolution, entailing neuroticism suppression and increased learning behaviors has been more significant in AMHs than in Neanderthals. However, adaptive evolution toward greater intelligence was more significant in Neanderthals than in AMHs. These findings suggest that mental traits involved in cultural development evolved differently in AMHs and Neanderthals, and that learning behaviors such as cooperative learning, rather than intelligence, played an important role in AMH evolution.

## Introduction

Anatomically modern humans (AMHs; *Homo sapiens*) emerged in Africa and expanded their habitat to almost every corner of the globe (1). While spreading out, AMHs developed increasing technological sophistication and adaptive knowledge, which promoted efficiency, enabling their survival in new environments (2). Presumably, the adaptation of mental and behavioral phenotypes would have occurred during this cumulative process of cultural evolution, prompting a global evolutionary trend based on the distribution of associated alleles. One such trend would have been adaptive evolution of the *ST8SIA2* gene (3, 4), associated with brain/neural function (5). Selective increases of its promoter types (haplotypes of three promoter single nucleotide polymorphisms [SNPs]), annotated as schizophrenia-associated non-risk types and showing lower promoter activity than the risk types, induced a trend toward lower promoter activity at the population level (population promoter activity: PPA) through out-migration from Africa of AMHs. This phenomenon can be interpreted as an adaptation to psychosocial stress induced during the process of adapting to surrounding social environments (3, 4) because psychosocial stress is the sole schizophrenia environmental risk factor dealt with by brain functions (6, 7) (Fig. S1).

Considering the above discussion, we applied evolutionary trends of schizophrenia-associated genetic loci showing selective increases of non-risk allelic types as yardsticks through the establishment of a new population functional score (PFS). Consequently, we detected global adaptive evolution of mental phenotypes involved in cumulative cultural evolution by examining the dominance of similar trends in their evolution (Fig. S2). To elucidate the significance of such adaptive evolution in AMHs, we further determined differences between phenotypic trends in AMHs and Neanderthals using a new population polygenic score (PPS). We subsequently examined positive selection on genetic loci associated with mental phenotypes showing such trend differences in Neanderthals.

## Results

### A yardstick PFS-26 for non-AFR

As a first step, we developed a PFS reflecting allelic function at the population level by focusing on the functional difference (*D*) between alleles. We could therefore apply the functional trend of the *ST8SIA2* gene as a yardstick in the analysis using alleles for which no functional information was available. Next, we examined the correlation in the PFS distributions of 26 populations (PFS-26s) with a global spread, encompassing Africa (AFR), Europe (EUR), South Asia (SAS), East/Southeast Asia (EAS), and the Americas (AMR), derived from the 1000 Genomes Project (8). Therefore, experimental estimation of *D* was unnecessary (see Materials and Methods; Figs. S2 and S3). The PFS-26 of the *ST8SIA2* gene (PFS-26_ST8SIA2_) was obtained by determining the difference in promoter activity referring to the ancestral risk type, which showed the highest activity (Fig. 1A). PPA and PFS were negatively correlated, and PFS-26_ST8SIA2_ showed a correlation coefficient (*ρ*) of −0.99 with the PPAs of the 26 populations, indicating the precise translation of PPA into PFS. Because PFS-26_ST8SIA2_ mainly reflects the selective increase of schizophrenia-associated non-risk types in non-AFR (3, 4) (see Fig. 1A), we used it as a yardstick for examining selection in non-AFR.

**Fig. 1.**
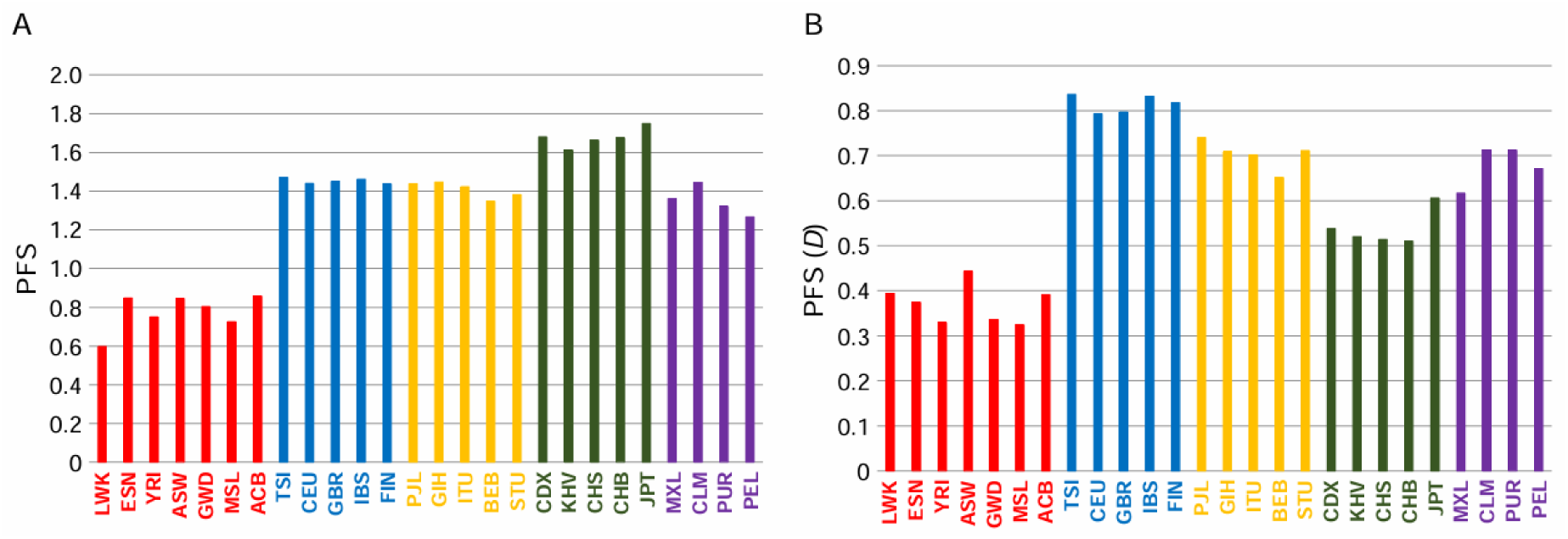
(A) PFSs of the promoter types of the *ST8SIA2* gene. (B) PFSs of the schizophrenia non-risk allele of rs1501357. Populations are represented by abbreviations used in the 1000 Genomes Project (8).

### A yardstick PFS-26 for AFR

To apply PFS-26 as a yardstick reflecting the selective increase of alleles in AFR, we next examined positive selection on schizophrenia-associated SNPs (9) (SCZ1 in Table 1). We detected signals of positive selection on non-risk alleles of three SNPs of brain/neural function-related genes implicated by fine-mapping (10): rs11693094 (*F*_c_ = 0.024, *P* = 0.003) in the *ZNF804A* gene, rs9922678 (*F*_c_ = 0.109, *P* = 4.3 × 10^−4^) in the *GRIN2A* gene, and rs1501357 (*F*_c_ = 0.028, *P* = 0.003) in the *HCN1* gene. The operational durations of this positive selection were estimated to be 17,100, 25,200 and 33,600 years for rs11693094, rs9922678, and rs1501357, respectively. These findings indicate that the detected positive selection could be an adaptation to psychosocial stress during the spread of AMHs. As for the non-risk alleles of rs11693094 and rs9922678, signals were detected in EUR and EAS, respectively. A signal on the non-risk allele of rs1501357 was also detected in AFR. We therefore selected its PFS-26 (PFS-26_rs1501357_) as a yardstick for examining positive selection in AFR (Fig. 1B).

**Table 1.**
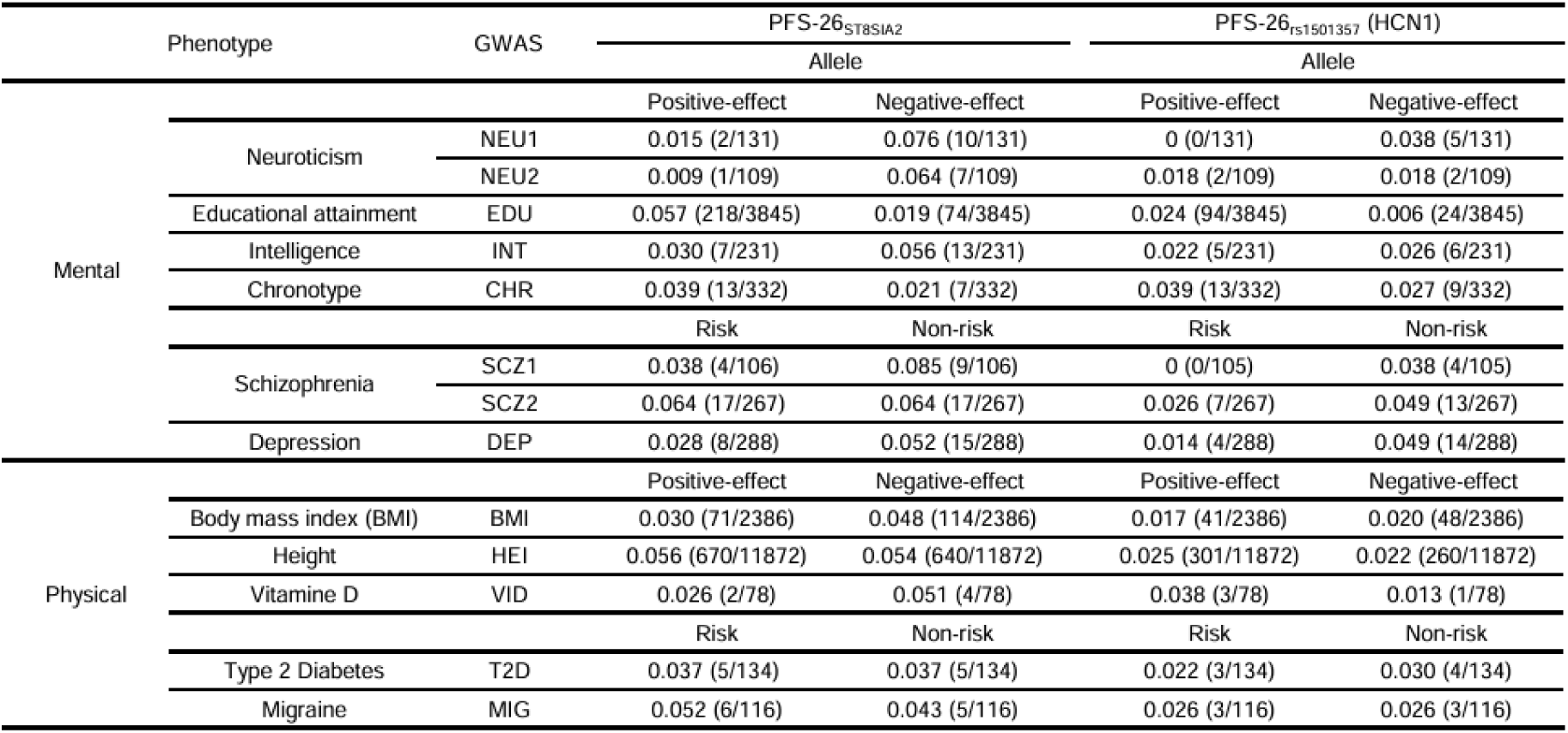
Observed rate of yardstick-like PFS-26s.

### Criterion in a correlation analysis

SNPs associated with phenotypes were obtained from genome-wide association studies (GWASs) (see Table 1 for abbreviations of the GWAS SNP sets), and index SNPs were therefore used instead of causal SNPs to obtain PFS-26s. Nearly complete linkages between index and causal SNPs can only be assumed in populations targeted in the GWAS analyses. Accordingly, the PFS-26s would not have been completely identical to those from causal SNPs. To assess the degree of discrepancy, we focused on concordant SNP pairs showing a tight linkage between the same allelic type of a particular phenotype in EUR because we obtained index SNPs from GWASs that mainly targeted the European population or meta-population containing Europeans (Fig. S4). The size of the associated region was typically <200 kb in length (11). In these regions, 743 concordant SNP pairs identified independently in different GWASs for the same phenotype (neuroticism, intelligence, schizophrenia, BMI, height, and type 2 diabetes; see Materials and Methods) showed a tight linkage (*r*^2^ > 0.81). Lastly, assuming that these tight-linked SNPs were index and causal SNPs, we examined correlations between PFS-26s in SNP pairs. The *ρ*-values were mainly distributed within a range from 0.9 to 1 (71% [527/743]) (Fig. S5A). Along a sliding window of *ρ* = 0.05, the PFS-26 proportion was consistently 6% (one-tenth of the highest proportion) or less after a window shift covering PFS-26s showing a *ρ*-value <0.9 (Fig. S5B). This indicated that when PFS-26 from a causal SNP was completely identical to the yardstick PFS-26s (i.e., *ρ* = 1), the PFS-26 from the index SNP generally showed a value of *ρ* > 0.9 in the correlation with the yardstick PFS-26. Thus, a *ρ*-value >0.9 was considered a reasonable criterion for assessing a high correlation with the yardstick PFS-26s.

The PFS-26s of non-risk alleles of rs11693094 and rs9922678, which showed positive selection in non-AFR (see above), exhibited a *ρ*-value >0.9 with PFS-26_ST8SIA2_ (*ρ* = 0.92 [rs11693094] and 0.92 [rs9922678]; see Fig. S6). The *ρ*-value of PFS-26_rs1501357_ with PFS-26_ST8SIA2_ was 0.68 (below the threshold of 0.9). These findings indicate that PFS-26_ST8SIA2_ reflects the PFS distribution resulting from selective allelic increase in non-AFR and are consistent with the assertion that PFS-26_ST8SIA2_ does not adequately document the selective increase of alleles in AFR. The criterion of *ρ* > 0.9 with the yardstick PFS-26s is therefore appropriate for detecting genetic loci showing a PFS distribution matching that of the yardstick PFS-26s. Furthermore, even if index SNPs are not guaranteed to be causal SNPs, this finding also suggests that index SNPs could be used instead of causal SNPs in our analysis.

### Convergent evolution of PFS-26s

The observed rate of PFS-26s exhibiting a *ρ*-value >0.9 with the yardstick PFS-26s (yardstick-like PFS-26s) was examined for each allelic type (Table 1). The rate for non-risk alleles/negative-effect alleles was compared with that for risk alleles/positive-effect alleles in each GWAS SNP set. Using PFS-26_ST8SIA2_, Bonferroni-corrected significant differences (α = 0.01/13; *P* < 7.7 × 10^−4^) were found in SNP sets associated with neuroticism (NEU1 and NEU2), educational attainment (EDU), and body mass index (BMI) (see Table 1), and the rates for negative-effect alleles of NEU1, NEU2, and BMI and positive-effect alleles of EDU were higher than those for the opposite-effect alleles (*P* = 3.7 × 10^−5^ [NEU1], *P* = 7.2 × 10^−5^ [NEU2], *P* = 1.1 × 10^−6^ [BMI], and *P* < 7.7 × 10^−4^ [EDU]; binomial test) (Table 1). The use of PFS-26_rs1501357_ showed the occurrence of Bonferroni-corrected significance in NEU1, EDU, SCZ1, and the depression-associated SNP set (DEP). The rates for negative-effect alleles of NEU1, positive-effect alleles of EDU, and non-risk alleles of SCZ1/DEP were higher than those for the opposite-effect alleles (*P* < 7.7 × 10^−4^ [NEU1, EDU and SCZ1] and *P* = 6.5 × 10^−5^ [DEP]; binomial test) (Table 1). The significant occurrence rate of the yardstick-like PFS-26s represents convergent evolution of PFS-26s through the global spread of AMHs (Fig. S2).

### The PFS-26s characteristic of phenotypes

To examine the involvement of the yardstick PFS-26s in PFS-26s unique to each phenotype, we obtained the PFS-26s characteristic of each allelic type showing a significant occurrence rate of the yardstick-like PFS-26s (see Fig. S7). We identified characteristic PFS-26s using the median *ρ*-value as a threshold to identify a similarity trend in a pairwise comparison of PFS-26s. These characteristic PFS-26s showed that *ρ*-values above the median value significantly occurred with the same allelic types rather than with the opposite-effect types (α = 0.05/number of alleles). They could be considered to reflect the unique allele distribution in a certain phenotype and were obtained in NEU1, NEU2, EDU, and BMI (Fig. 2 and Fig. S8). Strikingly, the distribution of *ρ*-values with the yardstick PFS-26s was restricted to a positive correlation in characteristic PFS-26s of negative-effect alleles of NEU1 (100% [8/8] in PFS-26_ST8SIA2_ and PFS-26_rs1501357_), NEU2 (100% [3/3] in PFS-26_ST8SIA2_), and BMI (100% [7/7] in PFS-26_ST8SIA2_), and positive-effect alleles of EDU (96% [2136/2223] in PFS-26_ST8SIA2_; 98% [2180/2223] in PFS-26_rs1501357_) (Fig. 2A–C and Fig. S8).

**Fig. 2.**
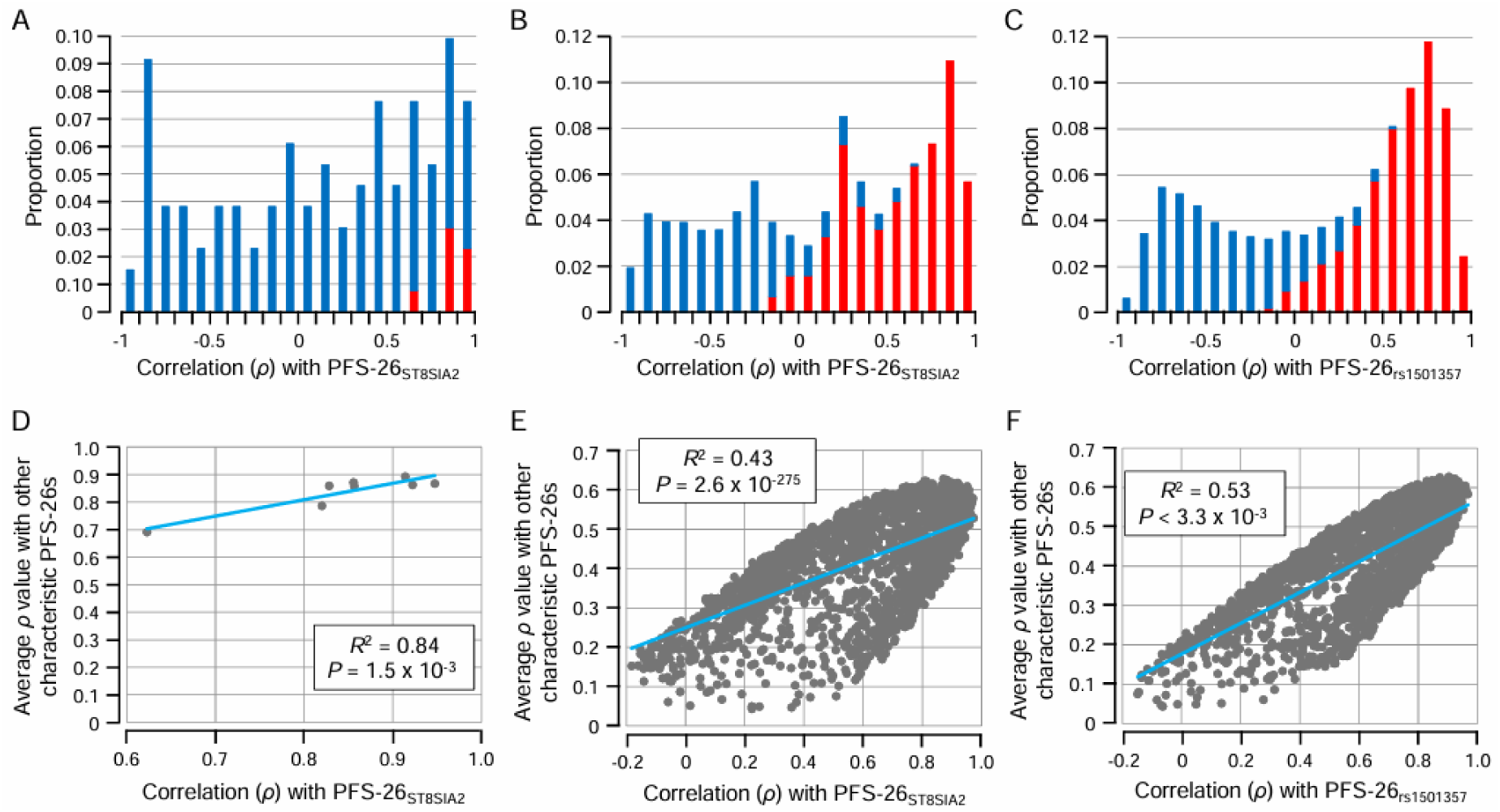
Involvement of the yardstick PFS-26s in the evolution of alleles associated with neuroticism and educational attainment. (A) Distribution of correlation coefficients with PFS-26_ST8SIA2_ in PFS-26s of negative-effect alleles of NEU1. (B) Distribution of correlation coefficients with PFS-26_ST8SIA2_ in PFS-26s of positive-effect alleles of EDU. (C) Distribution of correlation coefficients with PFS-26_rs1501357_ in PFS-26s of positive-effect alleles of EDU. The red part of each bar corresponds to the characteristic PFS-26s. (D) Correlation trend with PFS-26_ST8SIA2_ in the characteristic PFS-26s of negative-effect alleles of NEU1. (E) Correlation trend with PFS-26_ST8SIA2_ in the characteristic PFS-26s of positive-effect alleles of EDU. (F) Correlation trend with PFS-26_rs1501357_ in the characteristic PFS-26s of positive-effect alleles of EDU.

### Domination of the yardstick PFS-26s

The degree to which positive correlations with the yardstick PFS-26s dominated in the characteristic PFS-26s was elucidated by determining the correlation in characteristic PFS-26s between *ρ*-values with the yardstick PFS-26s and the average *ρ*-values with other characteristic PFS-26s (Fig. 2D–F; Fig. S9). As for PFS-26_ST8SIA2_, high *R*^2^ values with Bonferroni-corrected significant positive correlations (α = 0.01/3; *P* < 3.3 × 10^−3^) were found in characteristic PFS-26s of NEU1 and EDU (Fig. 2D and E). Higher *ρ*-values with PFS-26_ST8SIA2_ were associated with higher average *ρ*-values with other characteristic PFS-26s in negative-effect alleles of NEU1 (*R*^2^ = 0.84, *P* = 1.5 × 10^−3^) and positive-effect alleles of EDU (*R*^2^ = 0.43, *P* = 2.6 × 10^−275^) (Fig. 2D and E). A high *R*^2^ value with Bonferroni-corrected significant positive correlation was found for PFS-26_rs1501357_ in EDU (Fig. 2F). Higher *ρ*-values with PFS-26_rs1501357_ were associated with higher average *ρ*-values with other characteristic PFS-26s in positive-effect alleles of EDU (*R*^2^ = 0.53, *P* < 3.3 × 10^−3^) (Fig. 2F). Taken together with the significant occurrence rate of the yardstick-like PFS-26s (Table 1) and an exclusive restriction toward positive *ρ* values with the yardstick PFS-26s in the characteristic PFS-26s (Fig. 2A–C), the observed association shown in Fig. 2D–F indicated that a trend of becoming similar to the yardstick PFS-26s dominated for the PFS-26s unique to the negative-effect alleles of NEU1 and the positive-effect alleles of EDU.

To confirm the uniqueness of the domination of the yardstick PFS-26s in NEU1 and EDU, we applied the PFS-26s of alleles already established as selective targets (rs4988235 in the *MCM6* gene, rs1800414 in the *OCA2* gene, and rs3827760 in the *EDAR* gene) (12–14) as yardsticks (control-yardstick PFS-26s). The control-yardstick PFS-26s showed no dominant trend of becoming similar to them in the characteristic PFS-26s of NEU1 and EDU (Table S1 and Fig. S10).

### Global adaptation in AMHs

The dominant trend of becoming similar to the yardstick PFS-26s was detected only in the PFS-26s of neuroticism negative-effect alleles and educational attainment positive-effect alleles (Table 1 and Fig. 2). Considered together with the genetic correlation between lower neuroticism and increased educational attainment (15), this dominant trend indicates that singular global adaptive evolution has driven the suppression of neuroticism while increasing behaviors with positive effects on educational attainment. Neuroticism is a personality trait associated with sensitivity to psychosocial stress (16), whereas educational attainment is a behavioral phenotype primarily determined by environmental factors. Because the progress of cooperative learning via smooth communication that is achieved by overcoming psychosocial stress has a positive effect on educational attainment, psychosocial stress is considered an influential environmental factor. Tolerance to psychosocial stress is thus a shared function between neuroticism negative-effect alleles, educational attainment positive-effect alleles, and schizophrenia non-risk alleles (see also Fig. S1). We therefore posit that the dominant trend of becoming similar to the yardstick PFS-26s in the PFS-26s of neuroticism negative-effect alleles and educational attainment positive-effect alleles indicates a polygenic trend toward higher frequencies of allelic types conferring tolerance to psychosocial stress through the global spread of AMHs. Simply put, it indicates global polygenic adaptation to psychosocial stress.

### Assessment of Neanderthal phenotypes

We further examined the significance of detected global adaptive evolution in AMHs by comparing phenotypic trends for AMHs and Neanderthals (Altai, Vindija, and Chagyrskaya). Accordingly, we developed a new PPS by summarizing a polygenic phenotypic trend at the population level. This was based conceptually on the polygenic risk score/polygenic score (PRS/PS) in the GWAS analyses (see Materials and Methods). As previously noted, SNPs used in this study were index SNPs, which are not necessarily causal SNPs. We therefore examined the conservation of linkages between AMHs and Neanderthals using the 743 concordant SNP pairs showing a strong linkage (*r*^2^ > 0.81) in EUR. Because Neanderthal genome sequences are unphased, we selected 726 SNP pairs in which both SNPs are homozygous in Altai whose genome coverage was higher than Vindija and Chagyrskaya. Of these pairs, 589 (81%) showed complete linkages in Altai, indicating that using index SNPs is appropriate even in Neanderthals.

In every GWAS, over 99% of genetic loci were found in Neanderthals (see Table S2). This finding indicates that genetic loci associated with phenotypes targeted in this study have been well-conserved between AMHs and Neanderthals, and the estimation of PPS was performed using almost all of the genetic loci identified in the GWASs. Effect sizes showed a biased distribution toward smaller values in every GWAS (Fig. S11). These results indicate that phenotype expression depends on the orchestration of genetic loci having small effect sizes. Considering them together with almost complete conservation of genetic loci, we concluded that exclusion of genetic loci not obtained in Neanderthals was not so significant in our PPS comparison of AMHs and Neanderthals. Furthermore, even if Neanderthals had their own effect size in each genetic loci, differences from those of AMHs would be negligible.

### Significant phenotypic trends in AMHs

Given the conservation of gene repertories between AMHs and Neanderthals and the biased distribution toward smaller effect sizes (Fig. S11), we assumed that gene repertories and effect sizes were similar for AMHs and Neanderthals. We compared phenotypic trends in AMHs and Neanderthals using the PPS of positive-effect alleles (PPSp) (Fig. 3A; Fig. S12). In this comparative analysis, we focused on non-disease phenotypes because global adaptive evolution was only detected in these phenotypes (see Table 1 and Fig. 2). As for Neanderthals, the PPSp of each GWAS was estimated as a possible distribution because only three individuals were available (Fig. 3A; see Materials and Methods). When comparing PPSp in AMHs and Neanderthals, we detected Bonferroni-corrected significance (*α* = 0.01/7; *P* < 1.4 × 10^−3^) in NEU2, EDU, BMI, and the intelligence-associated SNPs (INT) but not in NEU1, the height-associated SNPs (HEI), and the chronotype-associated SNPs (CHR) (Fig. 3A; Fig. S12).

**Fig. 3.**
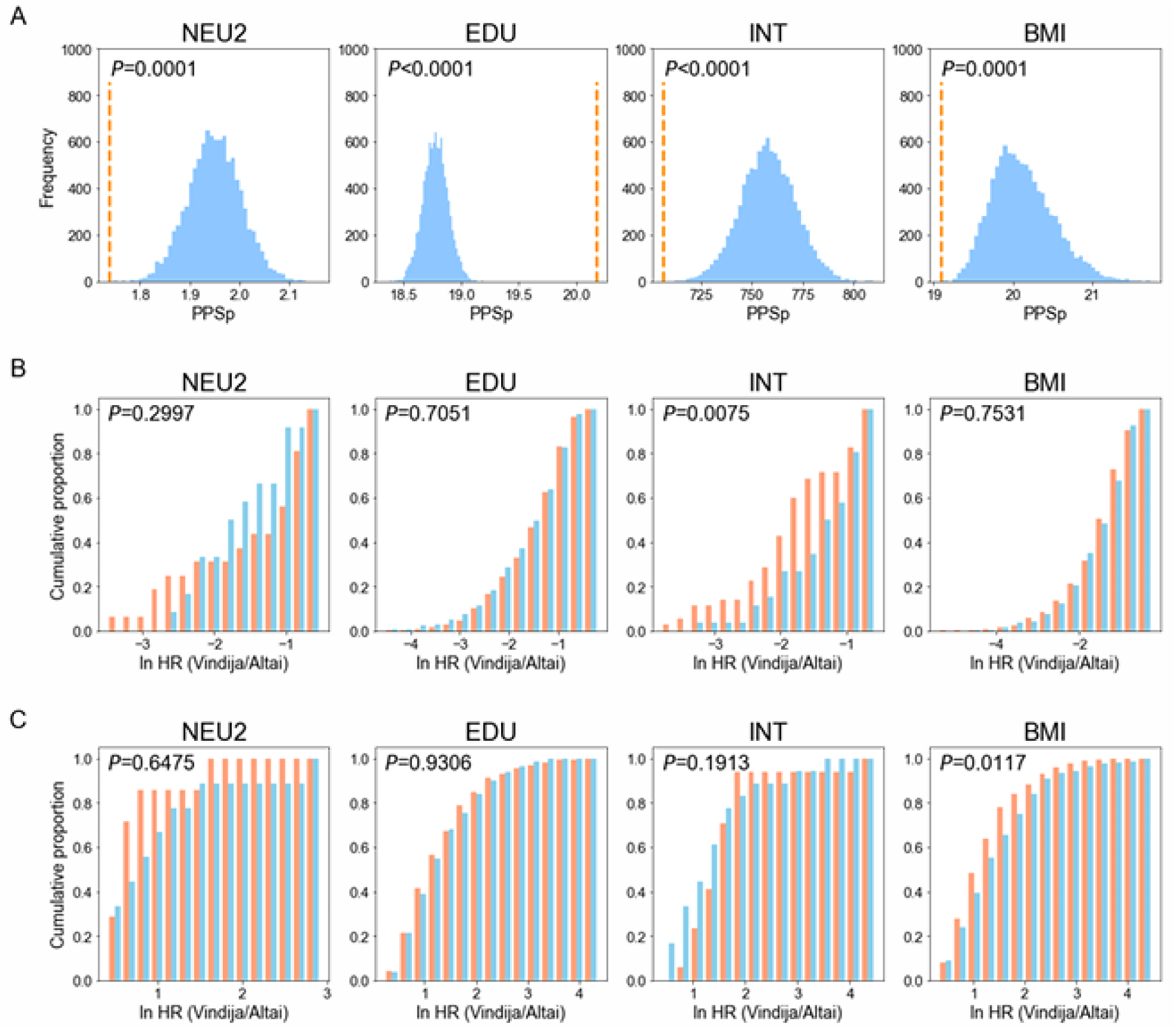
Phenotypic trends in Neanderthals. (A) A comparison of PPSp in AMHs and Neanderthals. The vertical dotted line shows a PPSp of AMHs. PPSp of Neanderthals is represented as its possible distribution. (B) Distribution of the heterozygosity ratio (HR) in the Vindija<Altai group. The red and blue bars represent positive-effect alleles and negative-effect alleles, respectively. (C) The distribution of HR in the Vindija>Altai group. Red and blue bars respectively represent positive-effect alleles and negative-effect alleles.

For BMI, PPSp was higher in Neanderthals than in AMHs (*P* = 1.0 × 10^−4^; Fig. 3A). This finding and the absence of a significant difference found in HEI (Fig. 3A; Fig. S12) were consistent with differences in body proportions between AMHs and Neanderthals, with Neanderthals being slightly shorter and significantly heavier than AMHs (17). These findings indicate that PPSp can be used to compare phenotypic trends of AMHs and Neanderthals.

For NEU2, the PPSp in AMHs was significantly lower than that in Neanderthals (*P* = 1.0 × 10^−4^; Fig. 3A). However, the PPSp for EDU was significantly higher in AMHs than in Neanderthals (*P* < 1.0 × 10^−4^; Fig. 3A). These findings are consistent with a view of global adaptive evolution toward the suppression of neuroticism and increased learning behaviors in AMHs. Given these consistencies in the two phenotypes, such global adaptive evolution is likely to have been more significant in AMHs than in Neanderthals. Thus, global adaptation to psychosocial stress was more significant in AMHs than in Neanderthals.

### Phenotypic trends in Neanderthals

Surprisingly, for INT, the PPSp of AMHs was significantly lower than that of Neanderthals (*P* < 1.0 × 10^−4^; Fig. 3A). Therefore, adaptive evolution toward greater intelligence was possibly more significant in Neanderthals than in AMHs. To assess this possibility, we also examined positive selection on genetic loci of NEU2, EDU, INT, and BMI in Neanderthals. Three Neanderthal individuals, Altai, Chagyrskaya, and Vindija, showed similar heterozygosity: 0.021%, 0.020%, and 0.019% (18). Because the ages of these individuals differed (Altai: 122,000 years; Chagyrskaya: 80,000 years; Vindija: 52,000 years) (18, 19), it can be assumed that heterozygosity was maintained constantly in the Neanderthal lineage. Considering the age difference of about 70,000 years between the oldest individual (Altai) and the youngest (Vindija), we examined the change in heterozygosity between them to look for positive selection (see Fig. S13). Our PFS analysis (see Table 1 and Fig. 2) revealed that a clear signal of positive selection was not necessarily detectable in all genetic loci identified by the GWAS. According to the difference in heterozygosity between Vindija and Altai (see Materials and Methods), the SNP sites, which are homozygous in both individuals, were categorized into three groups: Vindija<Altai (lower in Vindija), Vindija≈Altai (no difference) and Vindija>Altai (higher in Vindija) (Figs. S13 and S14). Positive selection at a particular site would reduce the heterozygosity of the region surrounding the site, and differences in heterozygosity between positive-effect alleles and negative-effect alleles would thus be observable, especially in the Vindija<Altai group and/or the Vindija>Altai group (Fig. S15). A significant difference (*P* < 0.05; Kolmogorov-Smirnov test) was detected in the Vindija<Altai group for INT and in the Vindija>Altai group for BMI (Fig. 3B and C).

The Vindija>Altai group for BMI showed that a trend toward lower heterozygosity in Vindija than in Altai was stronger in the positive-effect alleles than in the negative-effect alleles (*P* = 0.0117; Fig. 3C; Fig. S16). The higher PPSp for BMI in Neanderthals compared with that in AMHs remained unchanged even when Vindija was excluded (Fig. 3A; Fig. S17A), indicating that the detected difference in PPSp did not depend only on Vindija. These findings support the view that adaptive evolution toward higher BMI was more significant in Neanderthals than in AMHs. This interpretation is consistent with adaptive acquisition of body proportion (i.e., high BMI) in the Neanderthal lineage (17).

For INT, the Vindija<Altai group revealed that a trend toward lower heterozygosity in Vindija than in Altai was stronger for the positive-effect alleles than for the negative-effect alleles (*P* = 0.0075; Fig. 3B; Fig. S16). This means that intelligence-associated genetic loci were selective targets in Neanderthal populations. Furthermore, the higher PPSp for INT in Neanderthals compared with that in AMHs remained unchanged even when Vindija was excluded (Fig. 3A; Fig. S17B). These findings support the view that adaptive evolution toward greater intelligence was more significant in Neanderthals than in AMHs. We considered this finding together with the greater significance of global adaptive evolution toward the suppression of neuroticism and increased learning behaviors in AMHs than in Neanderthals. Accordingly, we posited that the mental traits associated with cultural development evolved differently in AMHs and Neanderthals (Fig. S18).

## Discussion

The yardstick PFS-26s (PFS-26_ST8SIA2_ and PFS-26_rs1501357_; Fig. 1) are just two PFS distribution patterns resulting from global adaptation to psychosocial stress. Given that other patterns would have been present but are currently unknown, the yardstick PFS-26s should not be misinterpreted as reflecting a difference in adaptation to psychosocial stress between populations.

NEU1 but not NEU2 showed a dominant trend of becoming similar to PFS-26_ST8SIA2_ (Table 1; Fig. 2A and D; Figs. S8A and S19A). However, NEU2 showed lower PPSp in AMHs than in Neanderthals, whereas NEU1 showed no significant difference in PPSp between AMHs and Neanderthals (Fig. 3A). In addition, for NEU1, the dominant trend of becoming similar to the yardstick PFS-26s was detected with the PFS-26_ST8SIA2_ but not with the PFS-26_rs1501357_ (Table 1; Fig. 2A and D; Figs. S8C and S19B). These incompatibilities relating to NEU1 and NEU2 could suggest a minim necessary locus size (∼100 SNPs) for our PFS and PPS analyses.

Although psychosocial stress is one of the environmental risk factors associated with the onset of schizophrenia and depression (6, 7, 20, 21), we did not detect any dominant trend toward becoming similar to the yardstick PFS-26s for SCZ1, SCZ2, or DEP. This could be attributed to the inclusion of their genetic loci associated with other environmental risk factors. The detection power of our analysis would have been reduced by the limited number of genetic loci associated with sensitivity to psychosocial stress. Importantly, this finding indicates that the yardstick-like PFS-26s did not result from negative selection against schizophrenia and depression.

Simply meeting the criterion of *ρ* > 0.9 with the yardstick PFS-26s is insufficient to warrant the conclusion that certain PFS-26s are involved in adaptation to psychosocial stress. To elucidate such involvement in each yardstick-like PFS-26 of NEU1, NEU2, and EDU, detection of positive selection would be needed, which is beyond the scope of this study.

The negative-effect alleles of BMI showed a significant occurrence rate of the yardstick-like PFS-26s (Table 1). This raises the possibility that adaptation to psychosocial stress also influenced physical phenotypes.

Henrich (2) argued that once individuals have evolved so that they can learn from each other with sufficient accuracy, social groups develop what could be termed ‘collective brains’, which drive cumulative cultural evolution. It is probable that overcoming psychosocial stress in the spread of AMH (i.e., global adaptation to psychosocial stress) expanded these collective brains by increasing the degree of social interaction and population size, thereby accelerating cumulative cultural evolution (4). We propose that the current genetic basis of neuroticism and educational attainment was established through such an evolutionary process (Fig. S20).

It appears that global adaptation to psychosocial stress was more significant in AMHs than in Neanderthals. This adaptation and its involvement in the enlargement of collective brains, leading to the evolution of collective brains, would have played a more important role in AMHs than in Neanderthals. This finding is consistent with larger population size and faster cultural evolution in AMHs than in Neanderthals (2, 18).

As noted above, the trend toward greater intelligence was plausibly more significant in Neanderthals than in AMHs. This trend and the greater significance of the collective brain in AMHs than in Neanderthals suggests that Neanderthals may have depended on functional improvements of individual brains rather than on the enlargement of collective brains for their survival. This could be related to larger average brain volume in Neanderthals (1,450 cc) than in AMHs (1,328 cc) (22). As Henrich (2) has argued, this difference could be related to the replacement of Neanderthals by AMHs.

In conclusion, the findings in this study suggest that mental traits (neuroticism, learning behaviors, and intelligence) associated with cultural development evolved differently in AMHs and Neanderthals, and that learning behaviors such as cooperative learning, rather than intelligence, played an important role in AMH evolution. We named ourselves *Homo sapiens* (wise man). The findings in this study may prompt a reconsideration of our self-recognition.

## Materials and Methods

### SNPs associated with phenotypes

Single-nucleotide polymorphisms (SNPs) associated with different phenotypes were obtained from genome-wide association studies (GWASs) reported to date (9, 10, 23–33) (see Table 1). SNPs identified according to genome-wide significance (*P* < 5.0 × 10^−8^) were used in this study.

Psychosocial stress is an environmental risk factor for the onset of schizophrenia and depressive disorders (6, 7, 20, 21). Neuroticism is also an environmental risk factor for the onset of conditions such as schizophrenia and depression (23), and individuals demonstrating higher neuroticism levels are vulnerable to social stress. We previously pointed to the possible role of adaptation to psychosocial stress in learning via communication (3, 4). Therefore, we selected GWASs of neuroticism (NEU1 and NEU2) (23, 24), educational attainment (EDU) (25), intelligence (INT) (26), schizophrenia (SCZ1 and SCZ2) (9, 10), and depressive disorder (DEP) (27) for inclusion in this study. In addition to educational attainment, we chose chronotype as a behavioral phenotype, and included a GWAS that examined it (CHR) (28). We also focused on GWASs conducted on physiological phenotypes (29–33): body mass index (BMI), height, (HEI), vitamin D level (VID), type 2 diabetes (T2D), and migraine (MIG) to examine whether any physiological factors contributed to adaptation to psychosocial stress.

Because the GWASs included those that did not apply conditional and joint multiple-SNP (COJO) analysis (34), we used SNP sets identified without COJO analysis, with the exceptions of HEI for which only SNP sets identified by COJO analysis were available. Referring to the database of the 1000 Genomes Project (hereinafter abbreviated as 1KGP) (8), we eliminated SNPs for which allelic frequencies were unavailable as well as SNPs located on the X chromosome and indels. For SNPs located on the X chromosome, the difference in scores (see below) among populations would have been inaccurate if the sex ratios of the populations differed. For example, the sex ratio for FIN in the 1KGP differed significantly from the expected 1:1 ratio of males to females (*P* = 0.02). In the GWASs used in this study, the SNPs on the X chromosome were as follows: 3 in SCZ1, 5 in SCZ2, 57 in EDU, 3 in CHR, 1 in VID, and 2 in MIG. Thus, they constituted a very small proportion of the SNPs in each GWAS. In this study, we mainly used GWASs that identified close to or more than 100 associated genetic loci to maintain statistical power for our analyses (see Tables S3–5). However, we also included GWASs that identified considerably fewer than 100 associated genetic loci (for neuroticism, intelligence, schizophrenia and type 2 diabetes) (35–43) in the case of linkage analysis using concordant SNP pairs (see the main text).

### DNA sequence data

The frequencies of each allele of the SNPs were retrieved from phase 3 of the 1KGP database (8), covering 2,504 individuals (5,008 sequences) from 26 different global populations. Referring to this database, we classified these populations into five meta-populations: Africa (AFR), Europe (EUR), South Asia (SAS), East and Southeast Asia (EAS), and the Americas (AMR).

DNA sequences of archaic humans, namely three Neanderthals (Vindija, Altai, and Chagyrskaya), were obtained from public databases: http://cdna.eva.mpg.de/neandertal/Vindija/; http://cdna.eva.mpg.de/neandertal/altai/; and http://ftp.eva.mpg.de/neandertal/Chagyrskaya/VCF/).

### Population functional score

Three SNPs (rs3759916, rs3759915, and rs3759914) in the promoter region of the *ST8SIA2* gene, which encodes a sialyltransferase synthesizing a linear homopolymer of sialic acid (polysialic acid) in the brain (5), are associated with schizophrenia risk (44–46). Selective increases of non-risk haplotypes of these three promoter SNPs (TCT and CGC types) occurred during the spread of AMHs (3, 4). These two promoter types show lower promoter activities than risk types, such as TGT and CGT types (3), resulting in a trend toward lower population promoter activity (PPA) in the spread of AMHs (4). This trend in the *ST8SIA2* gene can be interpreted as an adaptation to psychosocial stress and has been established in populations worldwide through the spread of AMHs (4). Thus, an examination of local adaptation is not sufficient on its own to identify similarity with this global functional trend in SNPs associated with phenotypes. Furthermore, because PPAs are estimated using both frequencies and the promoter activities of promoter types, they cannot be estimated for GWAS SNPs, whose functions are usually unknown. By focusing on differences in allelic function, we established a score that enabled us to examine the similarity of any SNPs to the global trend of population function of the *ST8SIA2* gene, even if the allelic functions of those SNPs were unknown. SNPs identified from GWAS analysis by their association with phenotypes are biallelic. By setting *D* as the difference in allelic function (e.g., promoter activity) in allele A1 from allele A2 at a biallelic SNP, and assuming additive allelic effects, the functional difference was expressed as 2*D*, *D*, and 0 for the A1/A1 homozygote, A1/A2 heterozygote, and A2/A2 homozygote, respectively. The functional difference at the population level (population function score: PFS) was obtained as follows:

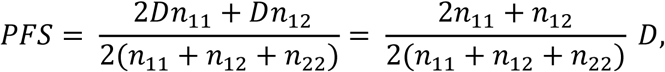

where *n*_11_, *n*_12_, and *n*_22_ are respectively the numbers of A1/A1 homozygotes, A1/A2 heterozygotes, and A2/A2 homozygotes in the population. Therefore, the PFS could also be defined simply as:

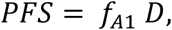

where 𝑓_*A*1_ is A1 allele frequency (2*n*_11_ + *n*_12_)⁄2(*n*_11_ + *n*_12_ + *n*_22_).

The PFS of the *ST8SIA2* gene was obtained using the actual functional difference (i.e., promoter activity). The TGT type is the ancestral type and shows the highest promoter activity (3). Thus, the PFS of the *ST8SIA2* gene (*PFS*_*ST*8*SIA*2_) was calculated as follows:

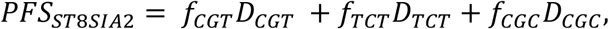

where *D_CGT_*, *D_TCT_*, and *D_CGC_* are differences in promoter activities from the TGT type in the CGT, TCT, and CGC types, and 𝑓*_CGT_*, 𝑓*_TCT_*, and 𝑓*_CGC_* are frequencies of the CGT, TCT, and CGC types, respectively.

Importantly, *D* of GWAS SNP was cancelled out in the estimation of the correlation coefficient (*ρ*) (i.e., similarity) between the distributions of PFSs from 26 populations (PFS-26; global trend of population function). Thus, correlation analysis using PFS-26 was performed to examine the similarity of the global trend of population function without requiring experimental estimation of allelic function (see Figs. S2 and S3).

In addition to eliminating the previously mentioned SNPs associated with phenotypes, we excluded those for which multiple derived alleles were indicated and those for which only derived alleles are found in AMHs. Table S3 shows numbers of SNPs used in the PFS analysis.

### Correlation with yardstick PFS-26s

The observed rate of high correlation with certain PFS-26s used as a yardstick was obtained for each allele type group (e.g., risk alleles of SCZ1). A binomial test was applied to examine whether this rate increased for certain allelic types compared with the opposite allelic types. The standard observed rate of PFS-26s showing a rate of high correlation with the yardstick PFS-26 is unknown in SNPs associated with phenotypes. We therefore compared the rate for negative-effect/non-risk alleles with that for positive-effect/risk alleles for each GWAS. Because populations show genetic kinship with one another, the assumption of independence among populations (see Fig. S3) does not hold when examining the correlation between PFS-26s. To enable strict assessment of the high correlation between PFS-26s, we needed a criterion that accounted for this genetic kinship (see the main text).

### Identification of characteristic PFS-26s

Similar PFS-26s uniquely found in a certain allelic type (e.g., schizophrenia risk alleles) were identified in a pairwise comparison of PFS-26s (Fig. S7). For their identification, we focused on the number of PFS-26s (*N*_>MED_) showing *ρ*-values exceeding the median value of all pairwise comparisons of PFS-26s. A particular type, of which *N*_>MED_ is significantly larger in comparing with the same allelic types than in that with the opposite-effect types (α = 0.05/number of alleles; χ^2^ test), is categorized as a characteristic PFS-26. In the χ^2^ test, results were not considered if *N*_>MED_ was <10.

### Detecting positive selection

We applied the *F*_c_ statistic to detect positive selection on SNPs in three meta-populations: AFR, EUR, and EAS (3, 47, 48). To detect all possible genetic loci, we modified a previously reported simulation method (3, 47) by including alleles showing an *S*-value (number of segregating sites) of at least 100. The Benjamini–Hochberg correction was performed for multiple testing. Significant discovery was then examined at a false discovery rate of < 0.1.

### Timing estimation on positive selection

We estimated the operational duration of positive selection following Fujito et al (3).

### Population polygenic score

Following the identification of SNPs associated with certain phenotypes in GWASs, the polygenic risk score (PRS)/polygenic score (PS) is generally applied to assess the degree of the phenotypic trend in individual (49). This score was obtained as follows:

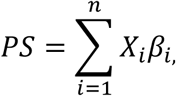

where *n* is the total number of associated SNPs, *X* is the number of alleles at associated SNPs in individuals, and *β* is the effect size estimated in GWAS. Accordingly, the population polygenic score (PPS) was developed to assess the phenotypic trend at the population level as follows:

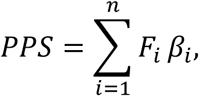

where *n* is total number of associated SNPs, *F* is the frequency of alleles at associated SNPs in a certain population, and *β* is the effect size estimated in GWASs.

### Estimation of PPSs in Neanderthals

Using Bayesian estimation based on allele frequencies in three Neanderthal individuals, we first obtained a possible distribution of the allele frequency at each genetic locus and subsequently applied this distribution to calculate the PPS. Lastly, we estimated the Neanderthal PPS in each GWAS as a distribution from 10,000 replications of the calculation. In addition to eliminating previously mentioned SNPs associated with phenotypes, we excluded SNPs not available in any Neanderthal individuals. The numbers of SNPs used in the PPS estimations are shown in Table S4.

### Classification of SNPs in Neanderthals

Homozygous SNP sites (i.e., positive-effect allele/positive-effect allele or negative-effect allele/negative-effect allele) in both Altai and Vindija were first selected as targets, and heterozygosity was calculated for the region available in both individuals by focusing on a 200-kb region (100-kb upstream and downstream) surrounding the target SNPs in AMHs. Target SNPs showing a 0 value for heterozygosity and those for which a 200-kb region was unavailable in AMHs were eliminated. As a final step, using the calculated heterozygosity value, we categorized target SNPs into three groups: Vindija<Altai, Vindija≈Altai, and Vindija>Altai according to the binomial distribution of heterozygosity for each SNP in Altai (see Figs. S13–S15). The Vindija<Altai group and the Vindija>Altai group contained genetic loci for which heterozygosity for Vindija was lower and higher than a 99% confidence interval (CI) of binomial distribution of heterozygosity for Altai, respectively. Genetic loci showing heterozygosity for Vindija within a 99% CI of heterozygosity for Altai were categorized in the Vindija≈Altai group. Table S5 shows the final number of SNPs used in the analysis.

## Data availability

The genomes used are available from the 1000 Genomes Project (phase 3 release, https://www.internationalgenome.org/).

## Acknowledgements

This research was supported by the Japanese Society for the Promotion of Science (grant nos. JP16K07535, JP19K06866, and JP24K09628 to T.H.). We thank Edanz (https://jp.edanz.com/ac) for editing a draft of this manuscript.

## Author contributions

T.H. conceived and supervised the study. Y.M., N.T.F., and T.H. collected sequence data. N.T.F., M.T., N.T., Y.S., and T.H. performed analysis detecting signals of positive selection. T.H. and Y.M. performed correlation analyses. Y.M. and T.H. performed analyses of phenotypic trends. Y.M., N.T.F., N.T., Y.S., and K.M.T. read and commented on the pre-final version of the manuscript. T.H. wrote the paper.

