## Supplementary Tables and Figures for "Evolution of mental traits linked to cultural development in modern humans and Neanderthals"

Supplementary materials include:
Figures S1 to S20
Tables S1 to S5

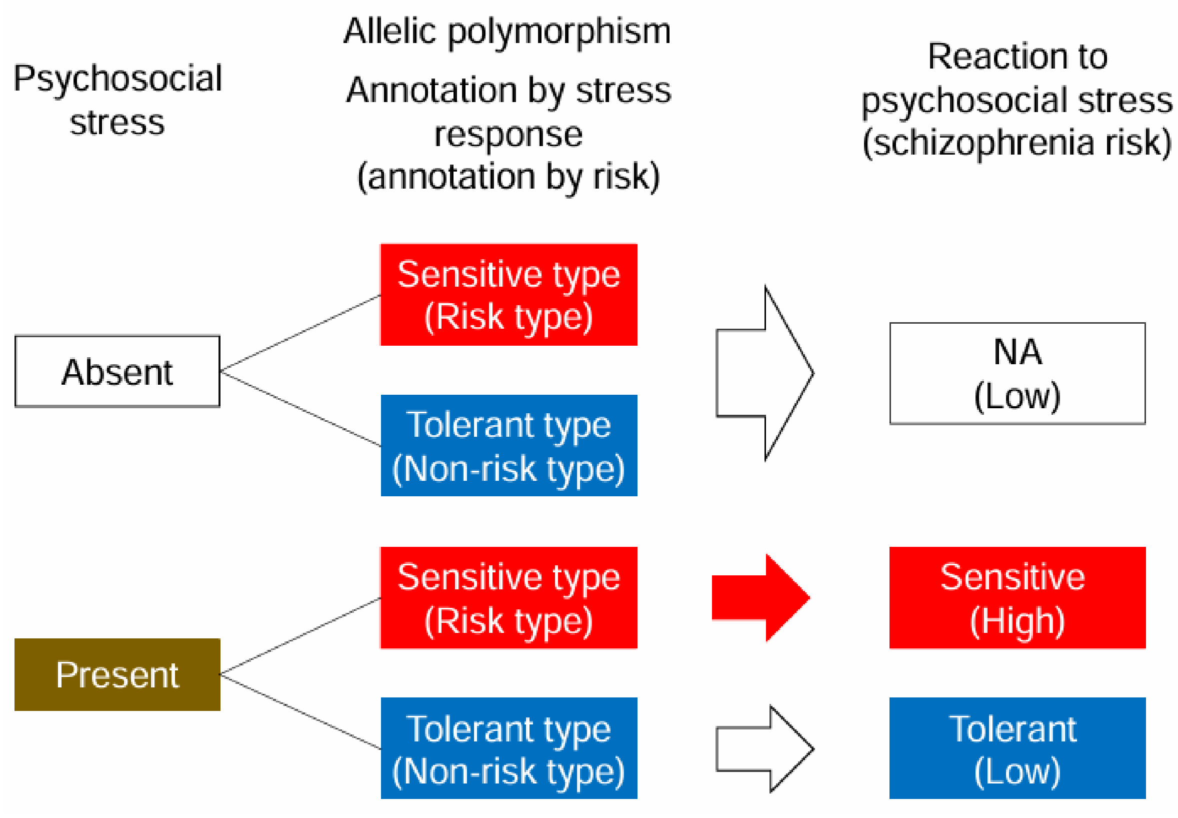


Fig. S1. The interaction between psychosocial stress and genetic polymorphism in schizophrenia onset. For simplicity's sake, only two cases of psychosocial stress (absent and present) are shown. Only in the presence of psychosocial stress do the non-risk type (tolerant type for psychosocial stress) and the risk type (sensitive type for psychosocial stress) differ functionally from each other. Positive selection on the non-risk (tolerant) type can therefore be interpreted as adaptation to psychosocial stress. NA=not applicable.

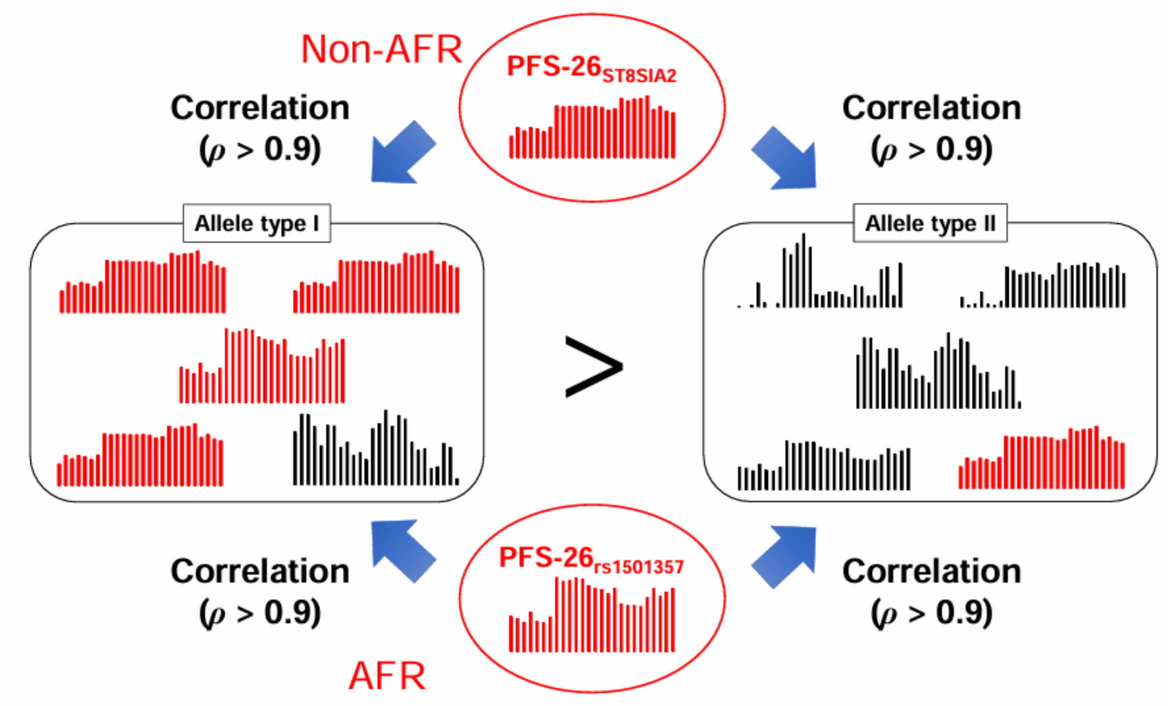


Fig. S2. The strategy for examining the convergent evolution of PFS-26s in this study.

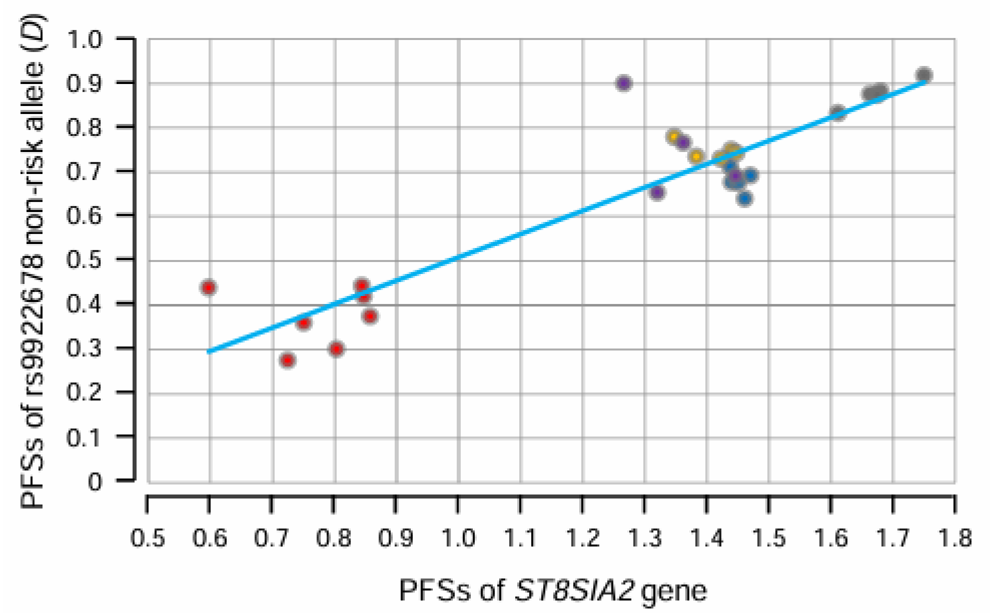


Fig. S3. A scatter plot of PFSs between the *ST8SIA2* gene and the non-risk allele of a schizophrenia-associated SNP (rs9922678; $\rho = 0.92$, $P = 3.7 \times 10^{-11}$). Three possible groups that reflect genetic kinship among populations were found in this plot, ruling out independence among populations.

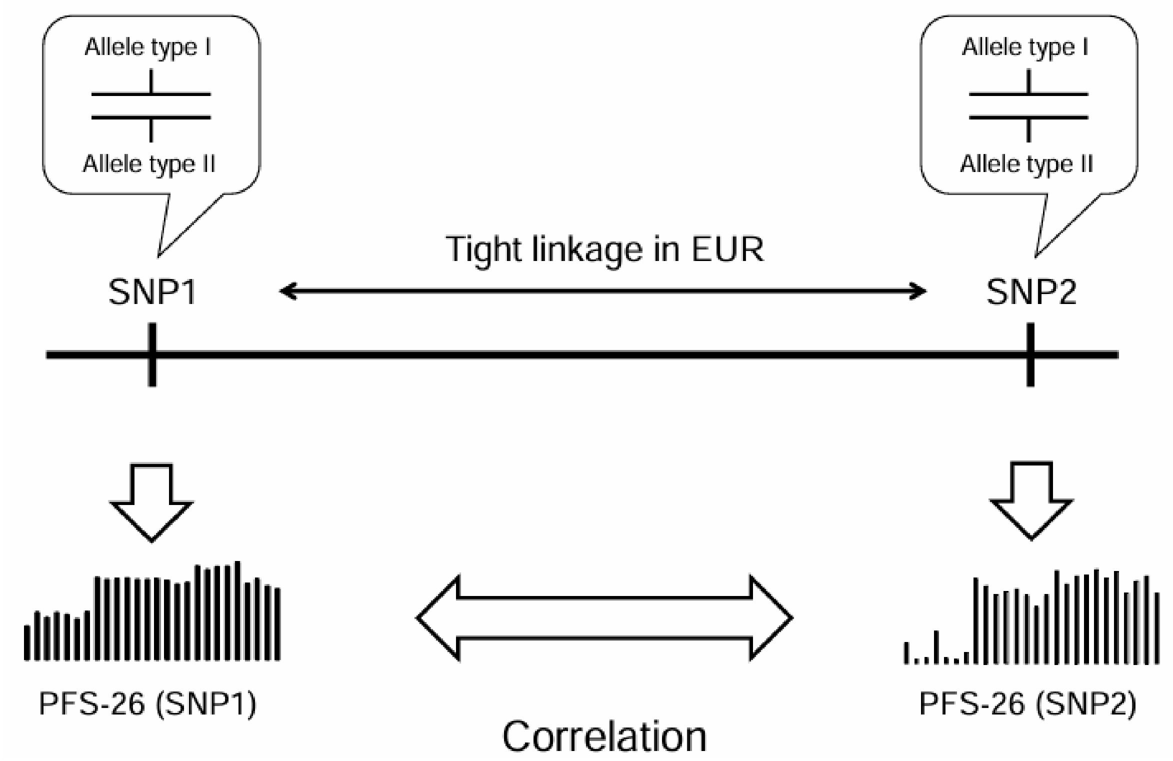


Fig. S4. The influence of linkage breakages in the PFS-26 correlation.

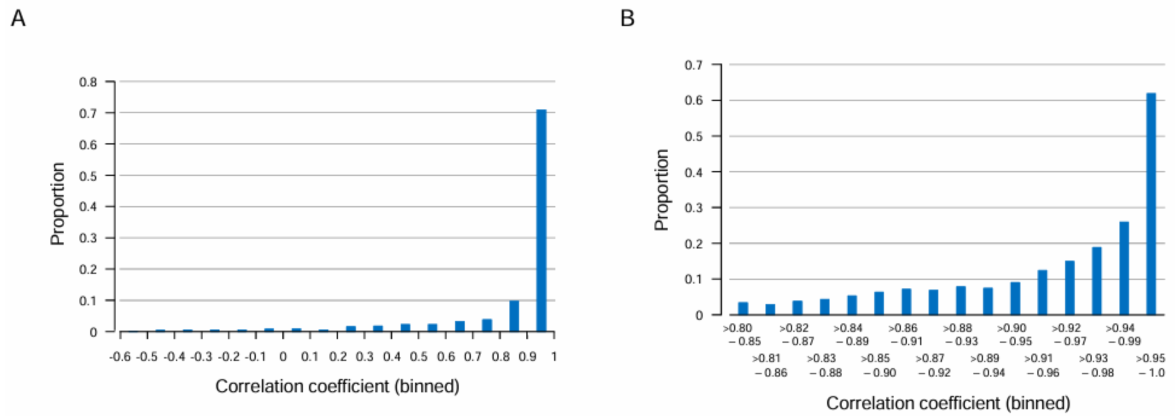


Fig. S5. Correlations between PFS-26s of tightly linked alleles in the European meta-population. (A) Distribution of correlation coefficients. (B) Distribution of correlation coefficients in the 0.8–1 range. Proportions were obtained using sliding windows of correlation coefficients (window size: 0.05; step size: 0.01).

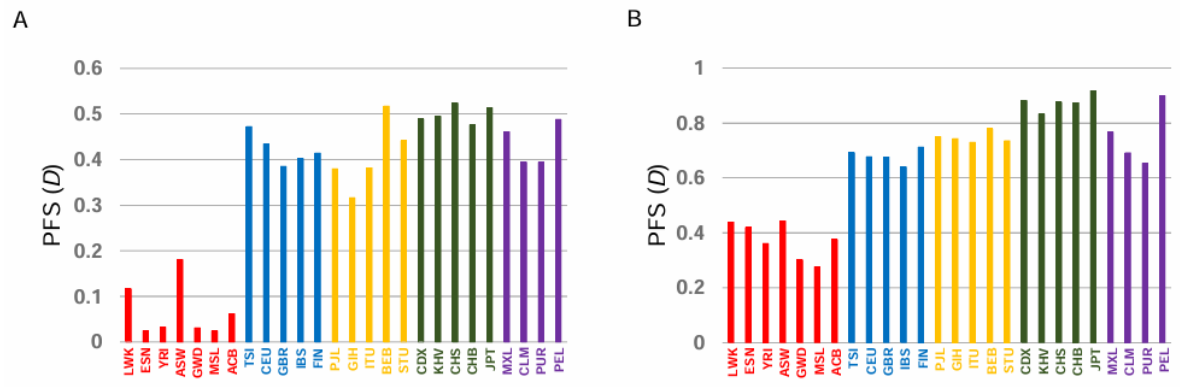


Fig. S6. PFS-26s of the non-risk alleles of rs11693094 (A) and rs9922678 (B).

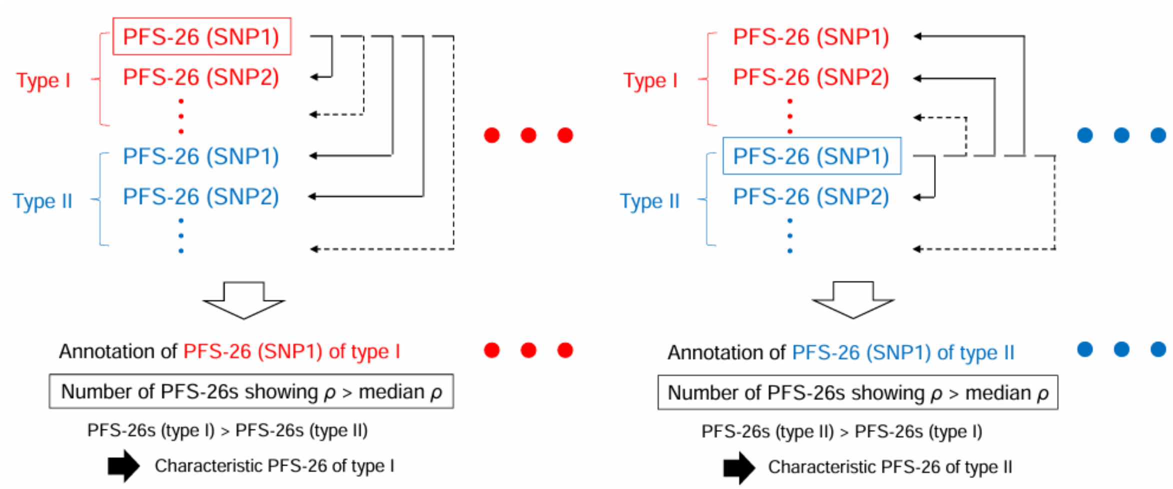


Fig. S7. Procedures for the identification of characteristic PFS-26s.

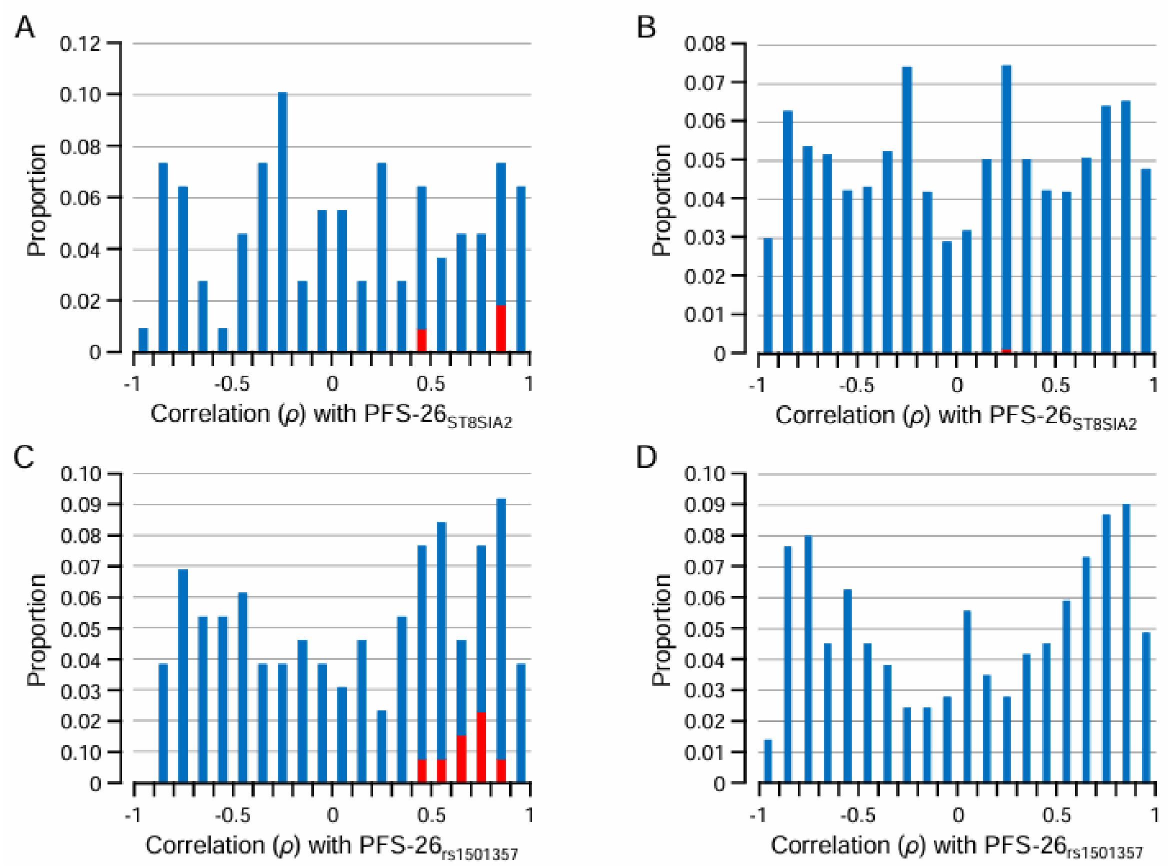


Fig. S8. Correlations with the yardstick PFS-26s in alleles associated with phenotypes. (A) Distribution of correlation coefficients with PFS-$26_{\mathrm{ST8SIA2}}$ in PFS-26s of negative-effect alleles of NEU2. (B) Distribution of correlation coefficients with PFS-$26_{\mathrm{ST8SIA2}}$ in PFS-26s of negative-effect alleles of BMI. (C) Distribution of correlation coefficients with PFS-$26_{\mathrm{rs1501357}}$ in PFS-26s of negative-effect alleles of NEU1. (D) Distribution of correlation coefficients with PFS-$26_{\mathrm{rs1501357}}$ in PFS-26s of non-risk alleles of DEP. The red portion of each bar corresponds to the characteristic PFS-26s.

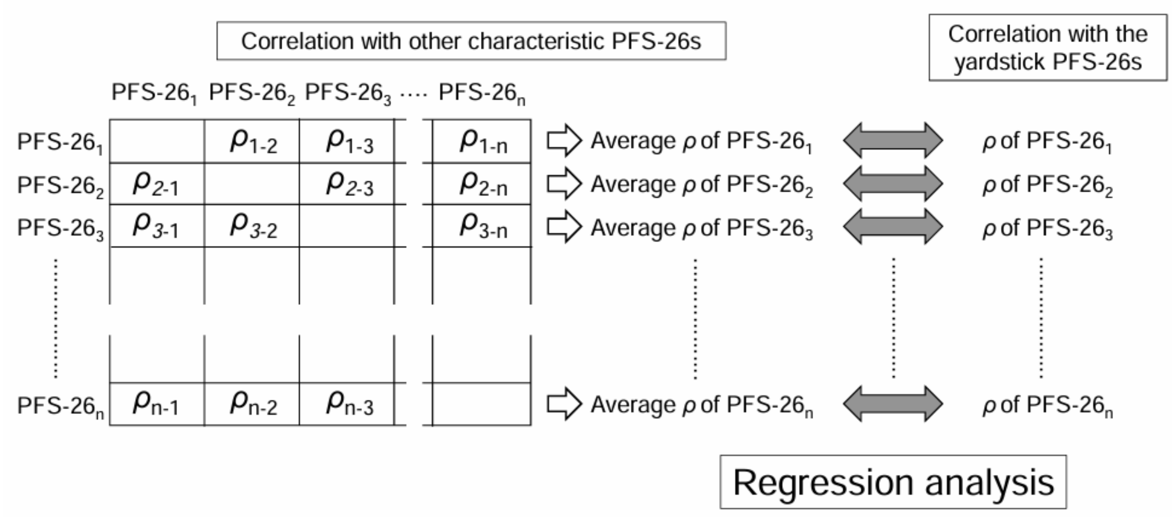


Fig. S9. Correlations in the characteristic PFS-26s between $\rho$-values with the yardstick PFS-26s and the average of $\rho$-values with other characteristic PFS-26s.

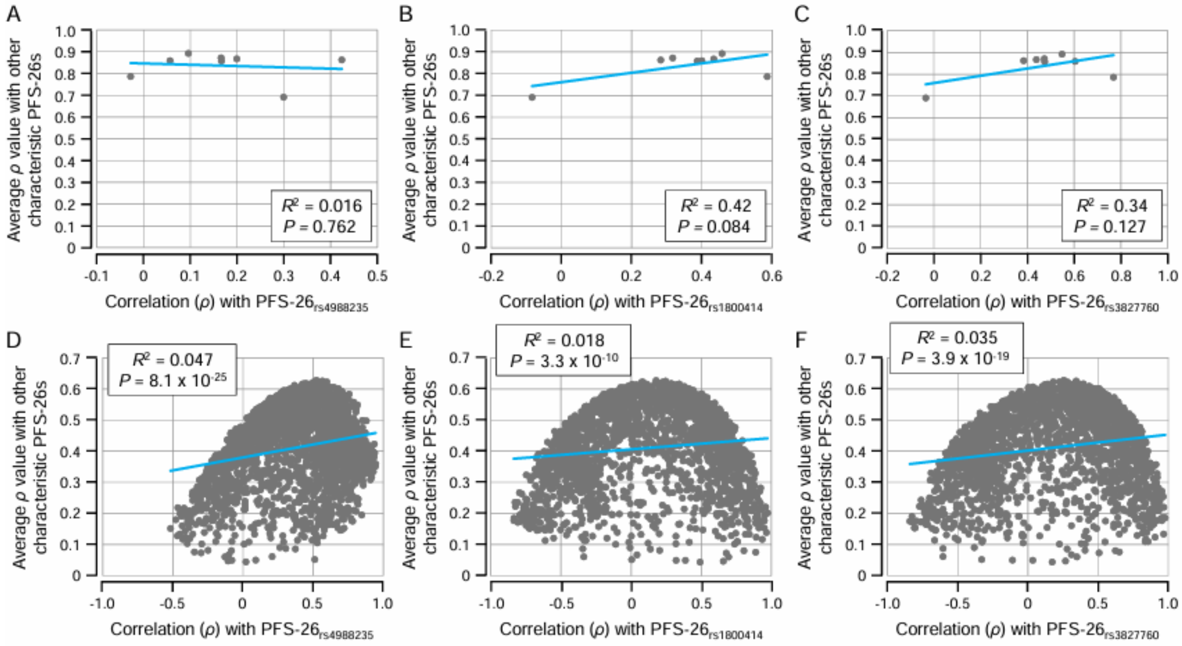


Fig. S10. Correlation trends with the control-yardstick PFS-26s in the characteristic PFS-26s of NEU1 and EDU. (A) PFS-$26_{rs4988235}$ in NEU1. (B) PFS-$26_{rs1800414}$ in NEU1. (C) PFS-$26_{rs3827760}$ in NEU1. (D) PFS-$26_{rs4988235}$ in EDU. (E) PFS-$26_{rs1800414}$ in EDU. (F) PFS-$26_{rs3827760}$ in EDU.

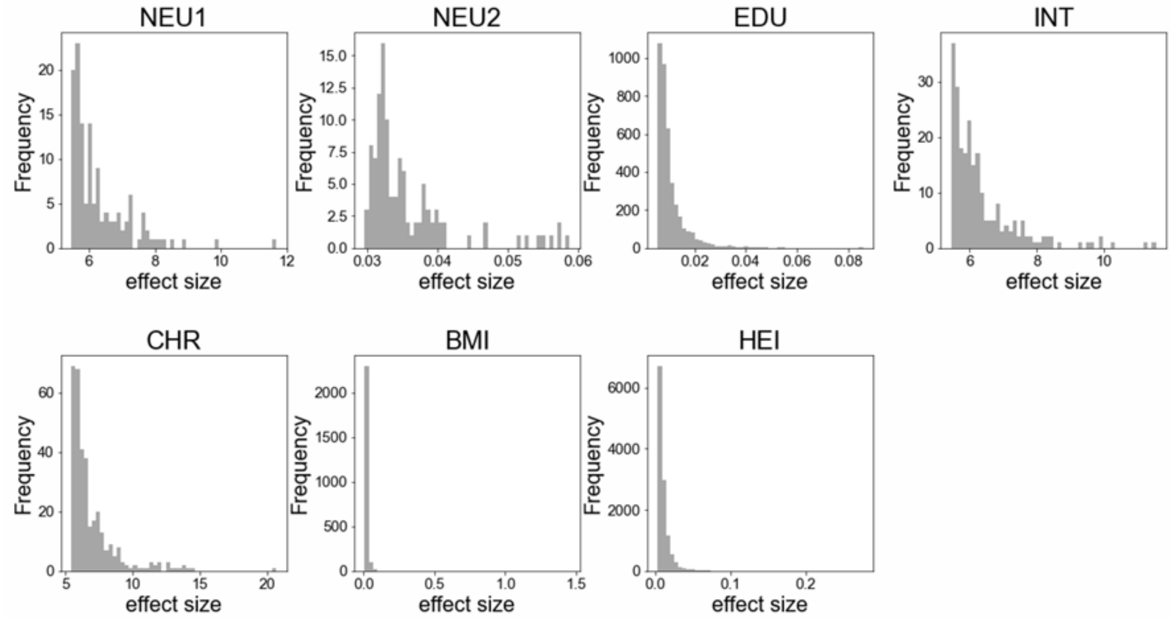


Fig. S11. Distribution of effect sizes of positive-effect alleles.

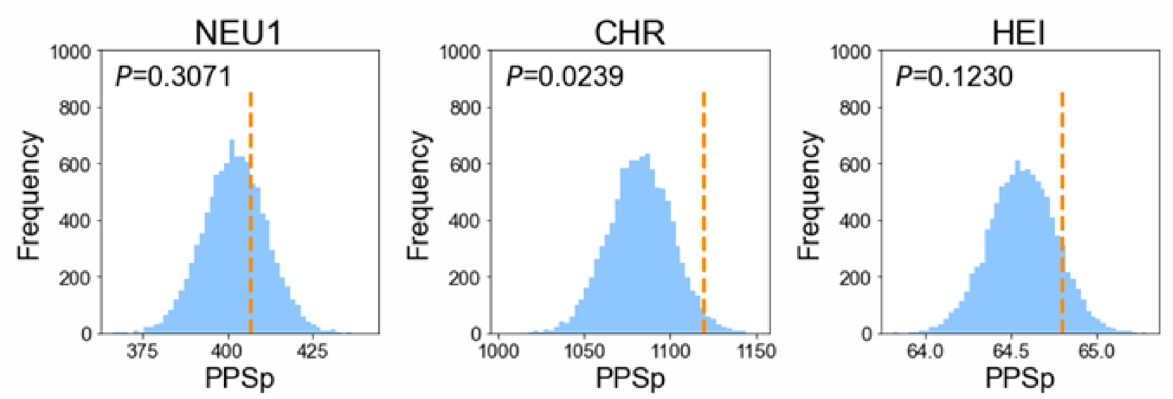


Fig. S12. A comparison of PPSp in AMHs and Neanderthals. The vertical dotted line shows the PPSp of AMHs. The PPSp of Neanderthals is represented as its possible distribution.

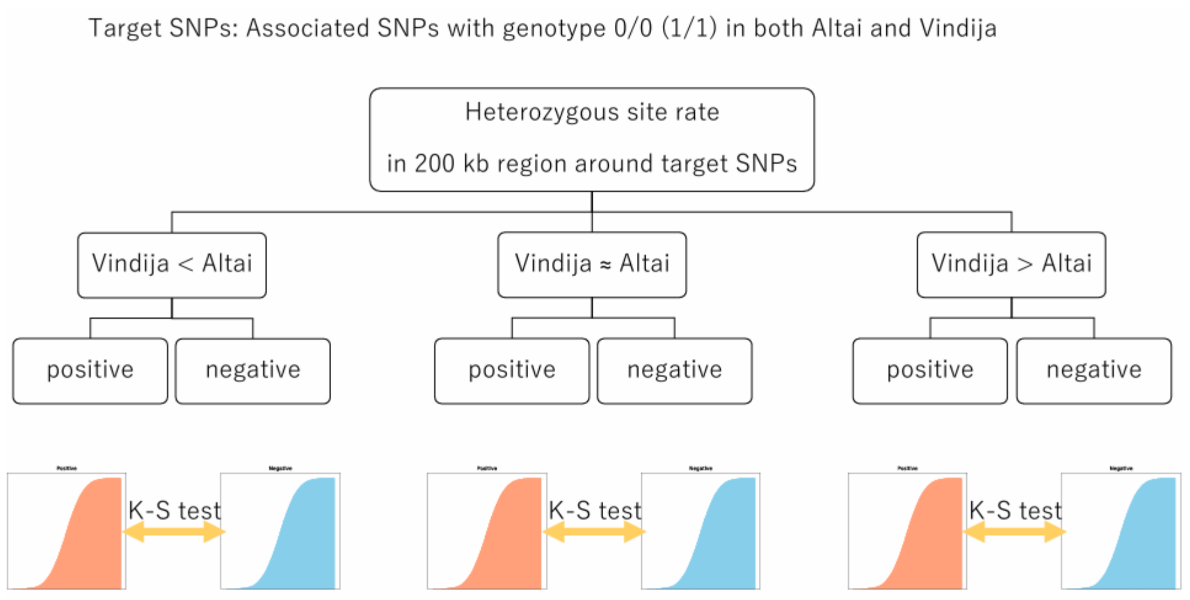


Fig. S13. The procedure used to detect positive selection in Neanderthals.

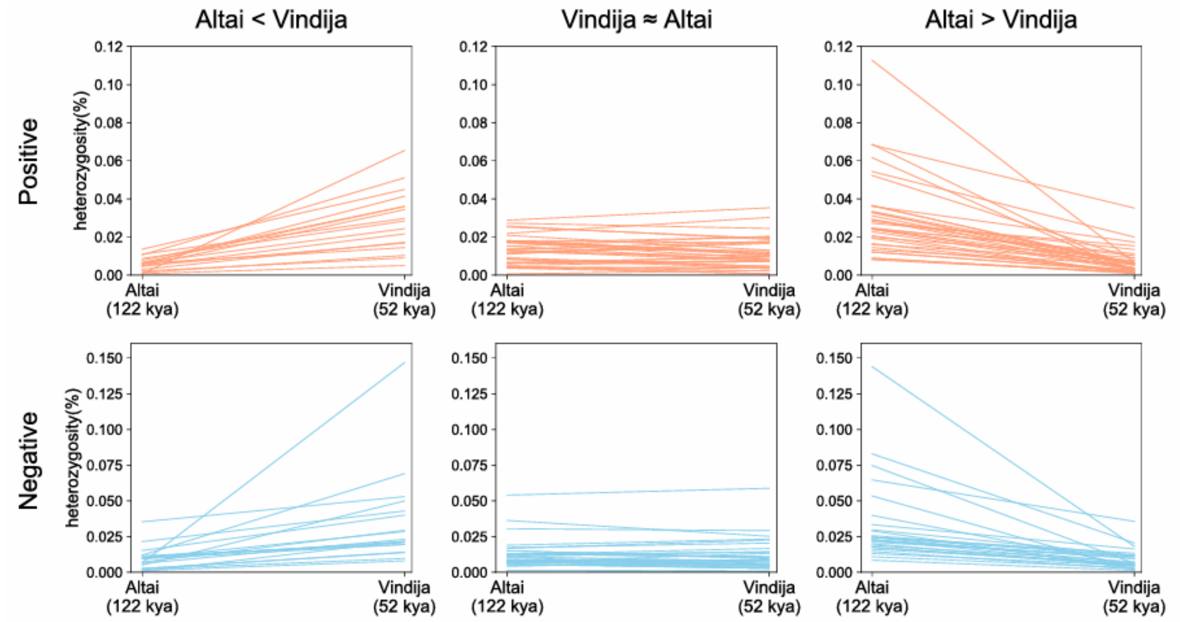


Fig. S14. Heterozygosity changes in the Vindija<Altai, Vindija≈Altai, and Vindija>Altai groups using the example of INT. kya, thousand years ago.

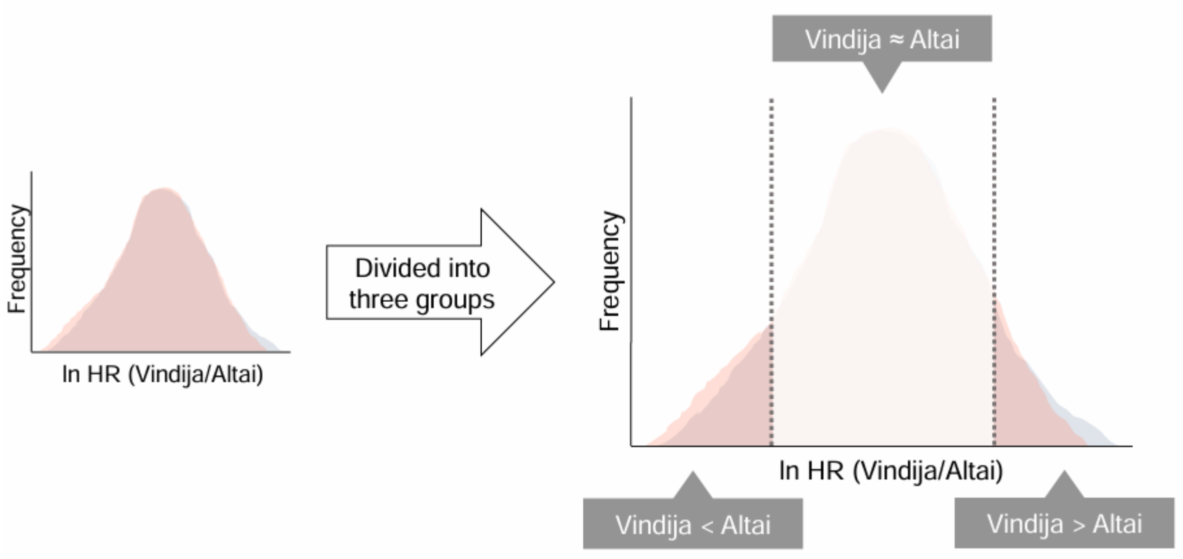


Fig. S15. Reduction in heterozygosity at genetic loci associated with phenotypes by positive selection. The distributions of the heterozygosity ratio (HR) in genetic loci for Altai and Vindija are displayed as overlapping positive-effect alleles (pink) and negative-effect alleles (blue). Given similar heterozygosity values for Altai (0.021%) and Vindija (0.019%) (18), the HR distribution of positive-effect alleles and negative-effect alleles would be similar under conditions of neutrality. However, if positive selection acted on positive-effect alleles, the HR distribution of positive-effect alleles would be reduced compared with that of negative-effect alleles. Accordingly, we divided SNPs into three groups: Vindija<Altai, Vindija≈Altai and Vindija>Altai groups (see Materials and Methods).

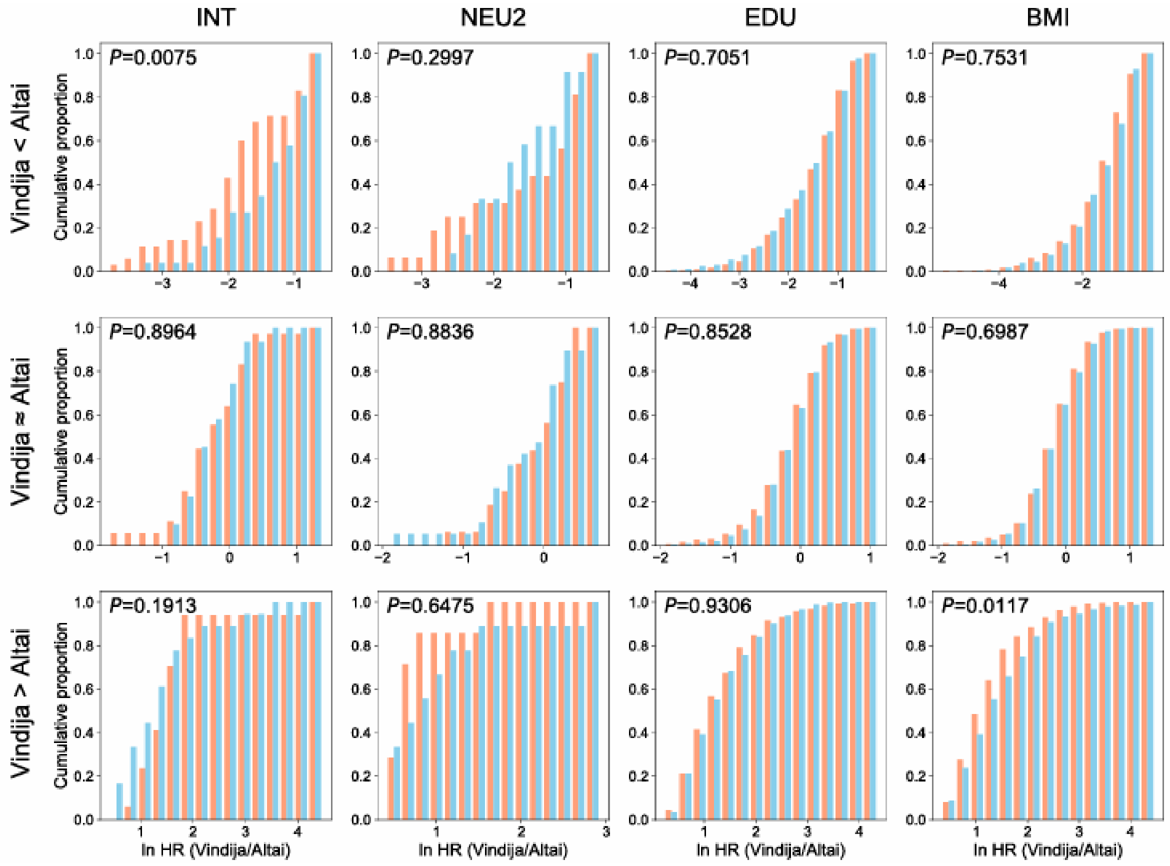


Fig. S16. Distribution of the heterozygosity ratio (HR) between Altai and Vindija. The red and blue bars respectively represent positive-effect alleles and negative-effect alleles. The Vindija<Altai and Vindija>Altai groups are shown in Fig. 3B and 3C.

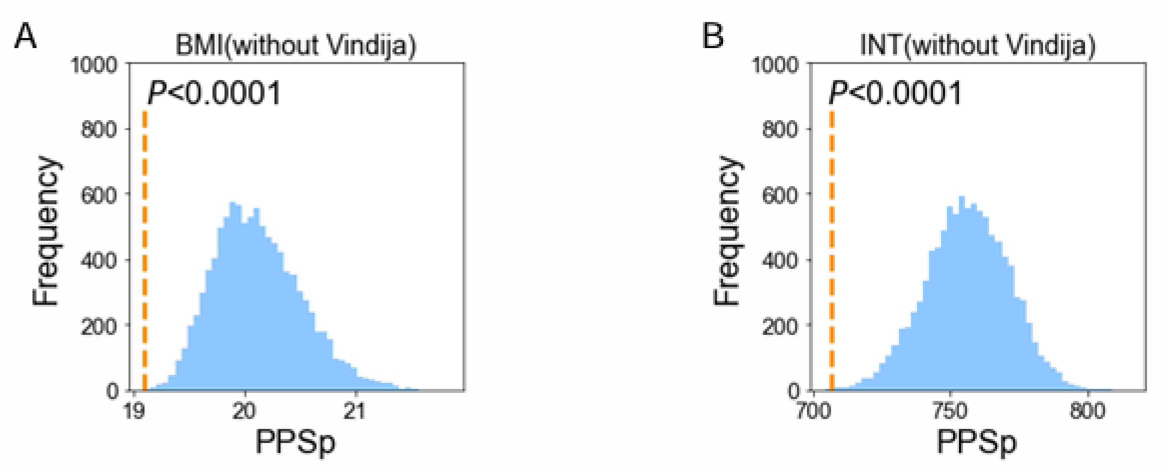


Fig. S17. Comparison of PPSp in AMHs and Neanderthals. (A) BMI. (B) INT. The vertical dotted line shows the PPSp of AMHs, while that of Neanderthals is represented as the possible distribution of PPSp. We obtained Neanderthals' PPSp distributions by excluding Vindija.

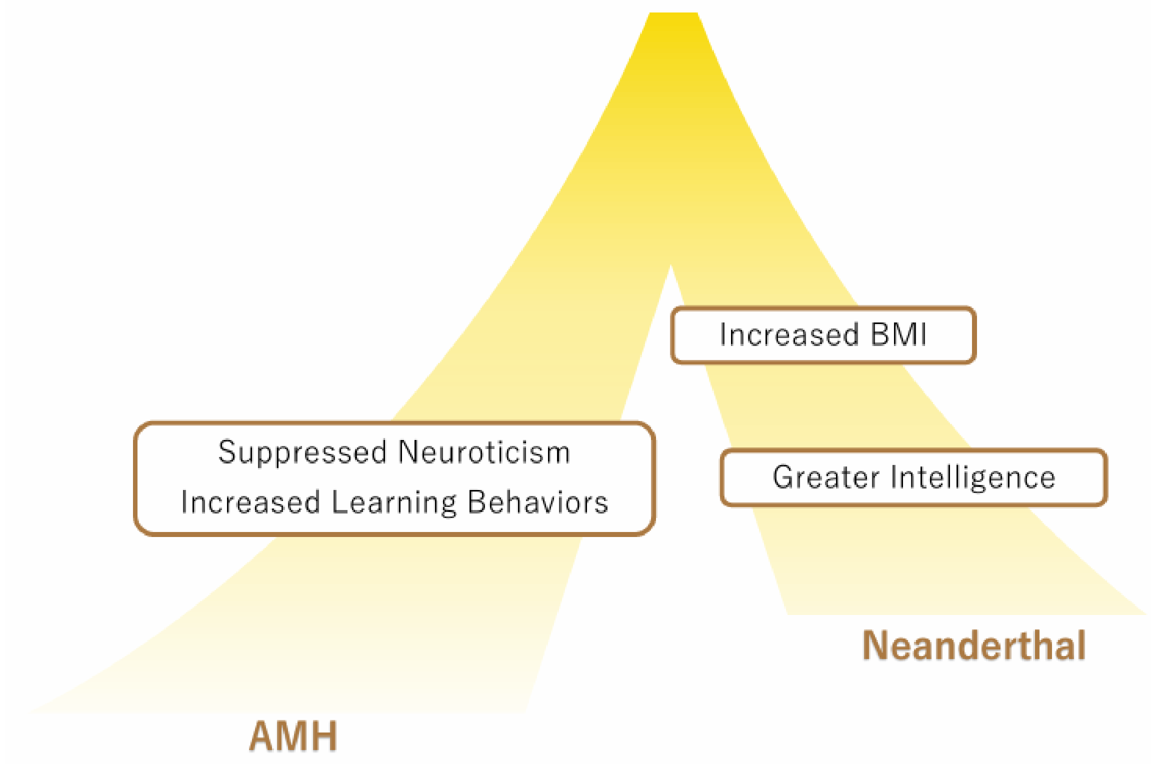


Fig. S18. Significant phenotypic trends in AMHs and Neanderthals.

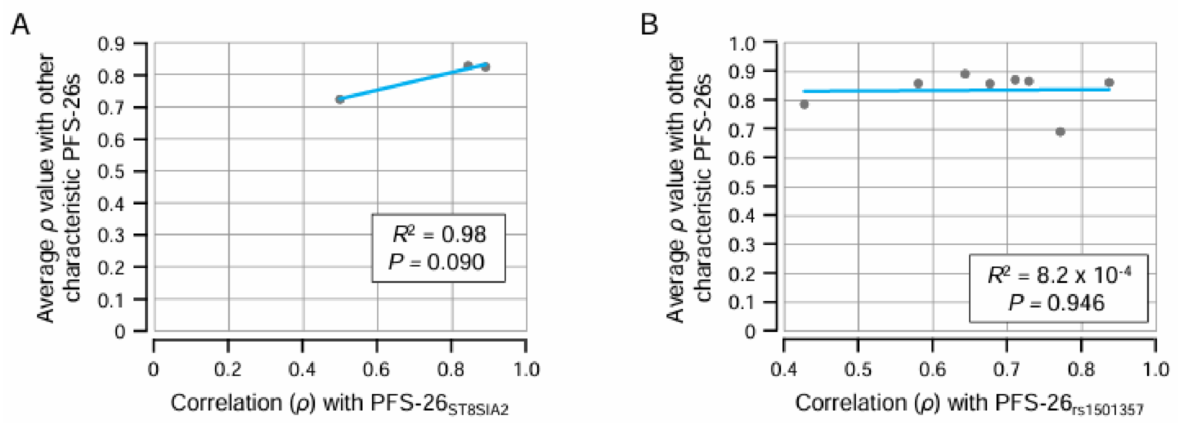


Fig. S19. The correlation trend with the yardstick PFS-26s in characteristic PFS-26s. (A) PFS-$26_{ST8SIA2}$ and the negative-effect alleles of NEU2. (B) PFS-$26_{rs1501357}$ and the negative-effect alleles of NEU1.

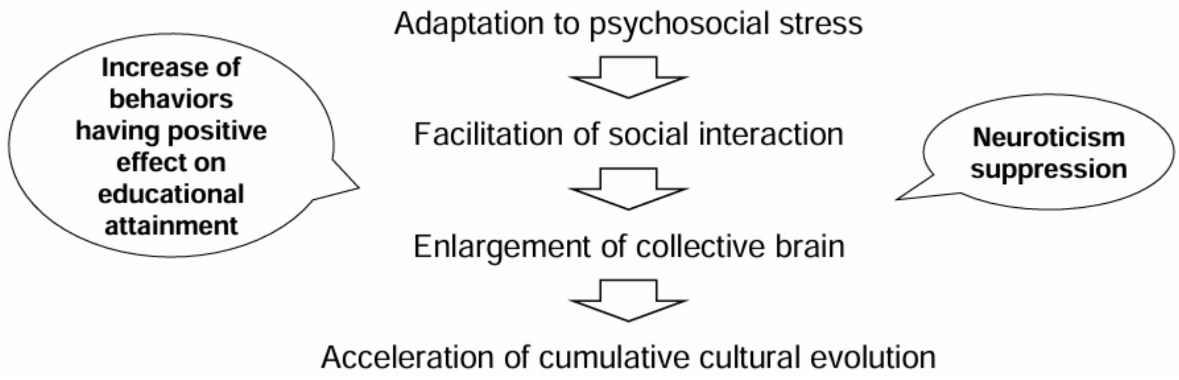


Fig. S20. A possible scenario depicting the role of adaptation to psychosocial stress in the evolution of anatomically modern humans.

Table S1. Observed rate of control yardstick-like PFS-26s.

| | Phenotype | GWAS | PFS-26$_{rs4988235}$ (MCM6) | | PFS-26$_{rs1800414}$ (OCA2) | | PFS-26$_{rs3827760}$ (EDAR) | |
|---|---|---|---|---|---|---|---|---|
| | | | Allele | | Allele | | Allele | |
| Mental | | | Positive-effect | Negative-effect | Positive-effect | Negative-effect | Positive-effect | Negative-effect |
| | Neuroticism | NEU1 | 0.008 (1/131) | 0 (0/131) | 0.008 (1/131) | 0.015 (2/131) | 0.015 (2/131) | 0 (0/131) |
| | | NEU2 | 0.009 (1/109) | 0.018 (2/109) | 0 (0/109) | 0.009 (1/109) | 0.009 (1/109) | 0.028 (3/109) |
| | Educational attainment | EDU | 0.006 (22/3845) | 0.003 (12/3845) | 0.007 (27/3845) | 0.004 (16/3845) | 0.010 (39/3845) | 0.006 (24/3845) |
| | Intelligence | INT | 0 (0/231) | 0.009 (2/231) | 0.009 (2/231) | 0.009 (2/231) | 0.004 (1/231) | 0.004 (1/231) |
| | Chronotype | CHR | 0 (0/332) | 0 (0/332) | 0.009 (3/332) | 0 (0/332) | 0.006 (2/332) | 0.006 (2/332) |
| | | | Risk | Non-risk | Risk | Non-risk | Risk | Non-risk |
| | Schizophrenia | SCZ1 | 0 (0/106) | 0 (0/106) | 0 (0/106) | 0 (0/106) | 0 (0/106) | 0.019 (2/106) |
| | | SCZ2 | 0.004 (1/267) | 0 (0/267) | 0.007 (2/267) | 0.007 (2/267) | 0.007 (2/267) | 0.011 (3/267) |
| | Depression | DEP | 0.003 (1/288) | 0 (0/288) | 0.003 (1/288) | 0.007 (2/288) | 0.024 (7/288) | 0.014 (4/288) |
| Physical | | | Positive-effect | Negative-effect | Positive-effect | Negative-effect | Positive-effect | Negative-effect |
| | Body mass index (BMI) | BMI | 0.006 (14/2386) | 0.004 (9/2386) | 0.008 (18/2386) | 0.006 (15/2386) | 0.011 (27/2386) | 0.008 (20/2386) |
| | Height | HEI | 0.001 (17/11872) | 0.001 (12/11872) | 0.011 (126/11872) | 0.009 (112/11872) | 0.013 (149/11872) | 0.012 (140/11872) |
| | Vitamine D | VID | 0 (0/78) | 0 (0/78) | 0.026 (2/78) | 0 (0/78) | 0 (0/78) | 0 (0/78) |
| | | | Risk | Non-risk | Risk | Non-risk | Risk | Non-risk |
| | Type 2 Diabetes | T2D | 0 (0/134) | 0 (0/134) | 0.007 (1/134) | 0.015 (2/134) | 0.007 (1/134) | 0.022 (3/134) |
| | Migraine | MIG | 0 (0/116) | 0 (0/116) | 0.009 (1/116) | 0 (0/116) | 0 (0/116) | 0 (0/116) |

Table S2. SNPs not obtained in Neanderthals

| | Phenotype | GWAS | SNP |
|---|---|---|---|
| Mental | Neuroticism | NEU1 | 0 |
| | | NEU2 | 0 |
| | Educational attainment | EDU | 1 |
| | Intelligence | INT | 0 |
| | Chronotype | CHR | 0 |
| Physical | Body mass index(BMI) | BMI | 1 |
| | Height | HEI | 5 |

Table S3. Phenotypes and GWASs used in the PFS analysis

| | Phenotype | GWAS | SNP* | | |
|---|---|---|---|---|---|
| Mental | | | Positive-effect | Negative-effect | Total |
| | Neuroticism | NEU1 | 68 | 63 | 131 |
| | | NEU2 | 57 | 52 | 109 |
| | Educational attainment | EDU | 2216 | 1629 | 3845 |
| | Intelligence | INT | 119 | 112 | 231 |
| | Chronotype | CHR | 184 | 148 | 332 |
| | | | Risk | Non-risk | Total |
| | Schizophrenia | SCZ1 | 45 | 61 | 106 |
| | | SCZ2 | 134 | 133 | 267 |
| | Depression | DEP | 139 | 149 | 288 |
| Physical | | | Positive-effect | Negative-effect | Total |
| | Body mass index (BMI) | BMI | 1170 | 1216 | 2386 |
| | Height | HEI | 6129 | 5743 | 11872 |
| | Vitamine D | VID | 39 | 39 | 78 |
| | | | Risk | Non-risk | Total |
| | Type 2 Diabetes | T2D | 70 | 64 | 134 |
| | Migraine | MIG | 61 | 55 | 116 |

* Annotation of derived alleles.

Table S4. SNPs used in the PPS analysis

| | Phenotype | GWAS | SNP |
|---|---|---|---|
| | | | Target SNPs |
| Mental | Neuroticism | NEU1 | 133 |
| | | NEU2 | 112 |
| | Educational attainment | EDU | 3912 |
| | Intelligence | INT | 232 |
| | Chronotype | CHR | 340 |
| Physical | Body mass index(BMI) | BMI | 2418 |
| | Height | HEI | 12051 |

Table S5. The number of SNPs used for the detection of positive selection in Neanderthals.

| | | Positive | | | | Negative | | | |
|---|---|---|---|---|---|---|---|---|---|
| | | Vindija < Altai | Vindija ≈ Altai | Vindija > Altai | Total | Vindija < Altai | Vindija ≈ Altai | Vindija > Altai | Total |
| Neuroticism | NEU2 | 16 | 16 | 7 | 39 | 12 | 19 | 9 | 40 |
| Educational attainment | EDU | 354 | 492 | 282 | 1128 | 493 | 643 | 411 | 1547 |
| Intelligence | INT | 35 | 36 | 17 | 88 | 26 | 31 | 18 | 75 |
| Body mass index(BMI) | BMI | 311 | 448 | 239 | 998 | 294 | 405 | 239 | 938 |
